# In-situ architecture of the human prohibitin complex

**DOI:** 10.1101/2024.02.14.579514

**Authors:** Felix Lange, Michael Ratz, Jan-Niklas Dohrke, Dirk Wenzel, Peter Ilgen, Dietmar Riedel, Stefan Jakobs

## Abstract

Prohibitins are a highly conserved family of proteins that have been implicated in a variety of functions including mitochondrial stress signalling and housekeeping, cell cycle progression, apoptosis, lifespan regulation and many others^1, 2^. The human prohibitins PHB1 and PHB2 have been proposed to act as scaffolds within the mitochondrial inner membrane, but their molecular organisation remained elusive. Using an integrative structural biology approach combining quantitative Western blotting, cryo-electron tomography, subtomogram averaging and molecular modelling, we determined the molecular organisation of the human prohibitin complex within the mitochondrial inner membrane. The proposed bell-shaped structure consists of eleven alternating PHB1 and PHB2 molecules. This study reveals an average of about 43 prohibitin complexes per crista, covering 1-3 % of the cristae membranes. These findings provide a structural basis for understanding the functional contributions of prohibitins to the integrity and spatial organisation of the mitochondrial inner membrane.

---

PHB1 (prohibitin or BAP32) and PHB2 (prohibitin-2, prohibitone, BAP37, REA) belong to the ancient and universally conserved SPFH (stomatin/prohibitin/flotillin/HflK/C) family of proteins^2^. A diverse set of critical functions has been attributed to the prohibitins. Although the localization of the prohibitins within the highly convoluted mitochondrial inner membrane is undisputed, additional cellular localizations are controversially discussed (Osman et al. 2009). Based on biochemical data and on negative-stain electron microscopy of prohibitin assemblies purified from yeast, it has been suggested that 12 to 20 PHB1/PHB2 heterodimers assemble into large ring complexes with a diameter of ∼20 nm^3, 4^. Such ring-like scaffolding structures might serve as scaffolds for proteins and lipids to form functional membrane domains. Still, direct experimental evidence for the existence, exact localization and number of ring-like prohibitin arrangements in mitochondria is missing.

First structural insights into assemblies formed by members of the SPFH family have been provided by studies expressing and purifying oligomers of the *E. coli* HflK/C proteins in complex with the client bacterial AAA+ protease FtsH. Subsequent single particle cryo-EM revealed that alternate HflK/C units assemble in a unique bell-shaped closed cage, containing 12 copies of each subunit^5, 6^. It remains unclear if this intriguing structure is a common feature also in-situ and if it is a unifying structural element of the entire SPFH family, or if the prohibitin sub-family of proteins developed a different, possibly a ring-like structure.

In this study, we aimed to visualize the putative prohibitin structure within the mitochondrial inner membrane by cryo-EM. Because of the conflicting reports on the sub-cellular localizations of the prohibitins, we first aimed at characterizing their localization in human U2OS cells^7, 8^. We found that transient expression of PHB1 or PHB2 fused to the fluorescent protein Dreiklang (DK) resulted in aberrant mitochondrial structures or cytoplasmic localization of the fusion protein, respectively (**Fig. S1A, B**)^9^. To explore if these effects are caused by the overexpression of the prohibitins or by their tagging with a fluorescent protein, we employed CRISPR-Cas9-mediated genome editing to generate human U2OS cell lines expressing PHB1 or PHB2 fused with DK from their respective native genomic loci, ensuring close to endogenous expression levels (**Fig. S2, S3, S4**). The mitochondrial network of both heterozygous cell lines (PHB1-DK and PHB2-DK) displayed wild type-like morphologies and PHB1-DK and PHB2-DK fusion proteins were localized to the mitochondria (**Fig. 1A, B**). This suggests that the overexpression, but not the tagging of the prohibitins with fluorescent proteins, induced artefacts.

**Fig. 1.:**
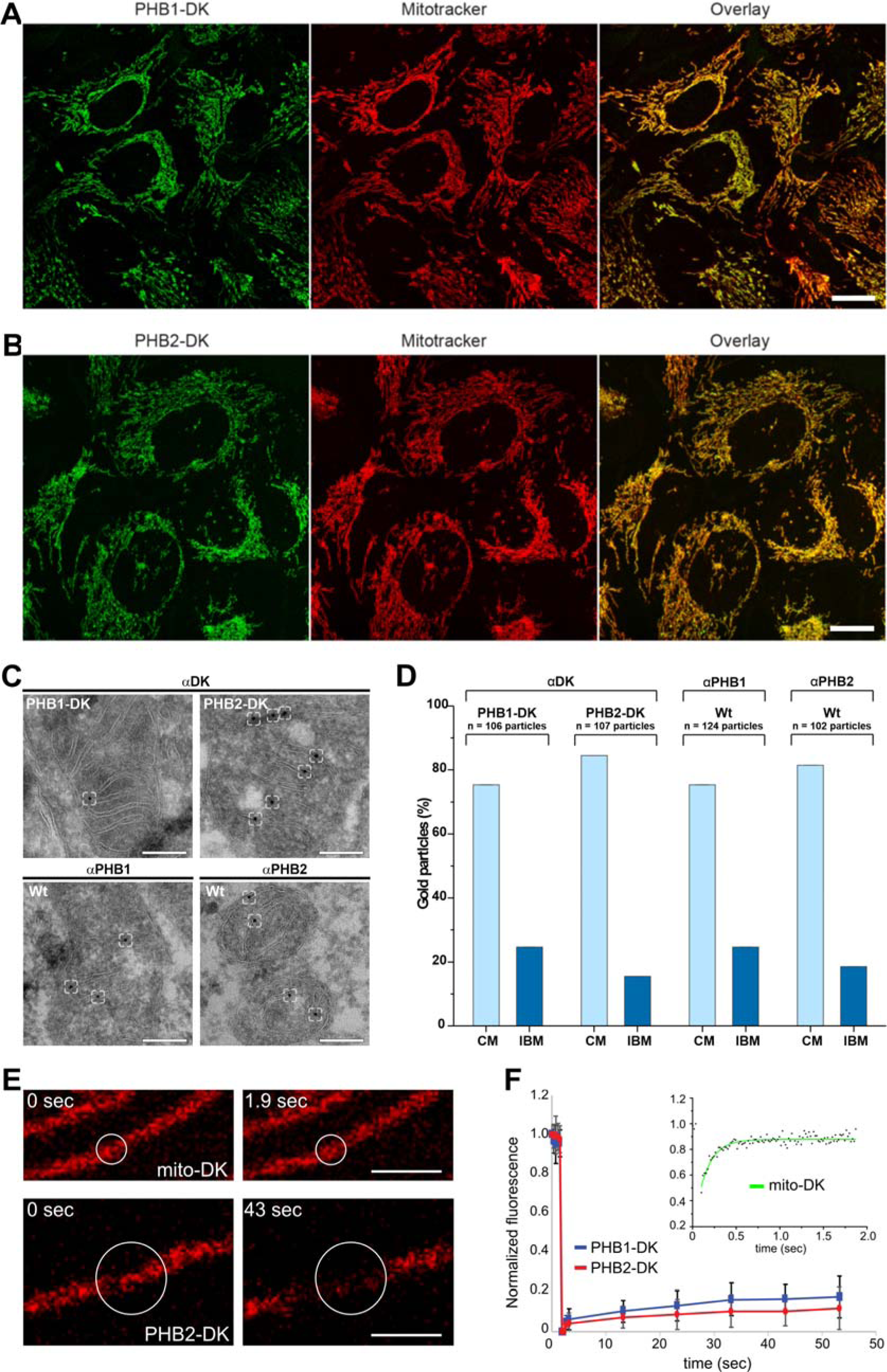
Human PHB1 and PHB2 are primarily localized at the crista membranes. **A, B:** Confocal microscopy of living cells expressing PHB1-DK (**A**) and PHB2-DK (**B**) at endogenous levels. Mitochondria were labelled using MitoTracker Deep Red FM (red). Dreiklang (DK) fluorescence is shown in green. **C:** Immunogold electron microscopy of PHB1-DK and PHB2-DK (top), and wild type (bottom) cells. Representative micrographs for each decoration with the indicated antibody are shown and the localization of gold particles is indicated by boxes. **D:** Quantification of the distribution of the gold particles. **E:** FRAP analysis of PHB2-DK compared to matrix-localized fluorescent protein Dreiklang (mito-DK). Circle represents the area before (0 sec) and after photobleaching in a representative experiment using mito-DK (1.9 sec) or PHB2-DK (43 sec), respectively. **F:** Recovery curves of PHB1-DK and PHB2-DK compared to mito-DK (inset). Scale bars: 10 µm (A, B), 200 nm (C), 2 µm (E).

**Fig. 2.:**
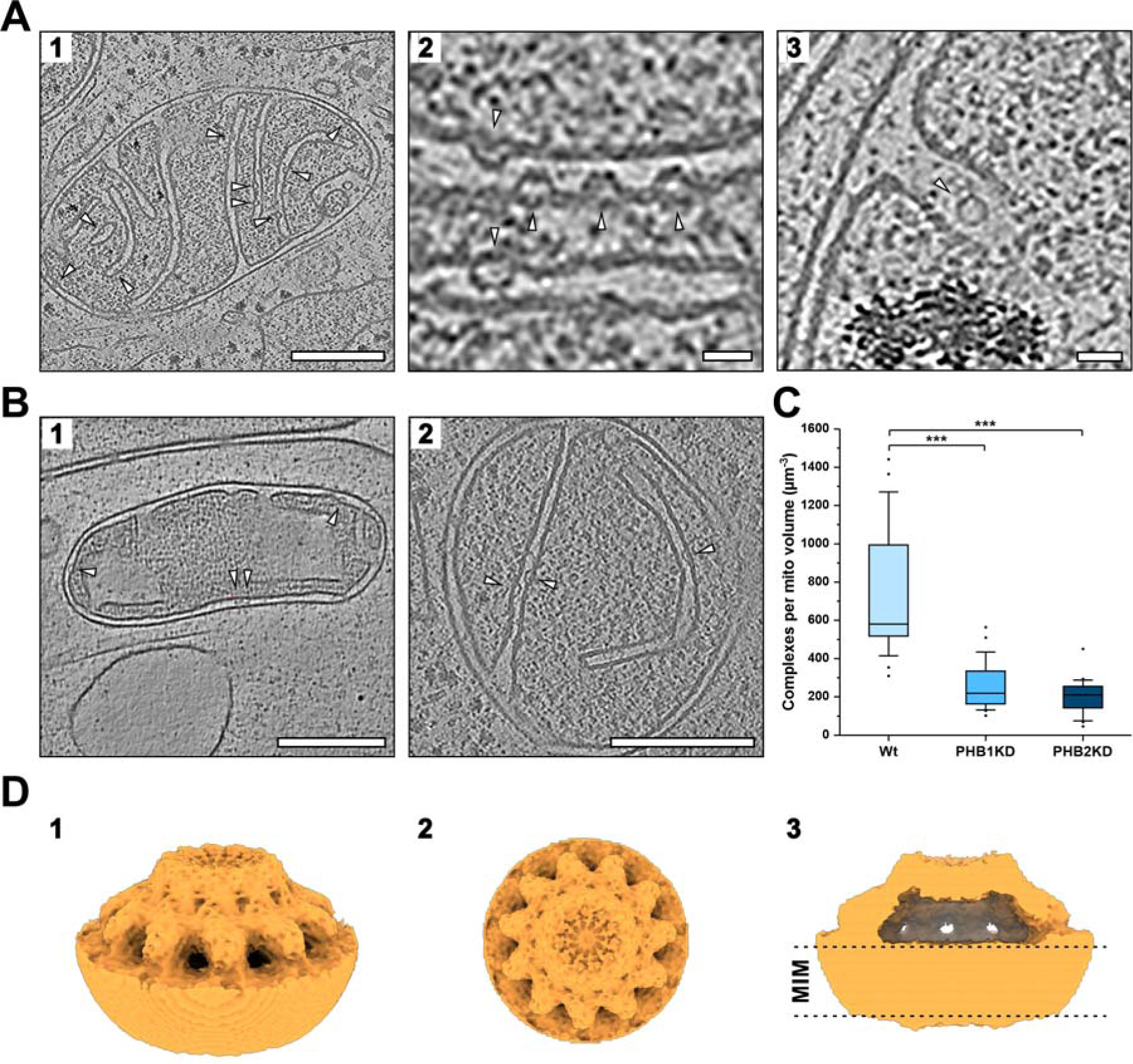
Prohibitins form bell-shaped complexes at the mitochondrial inner membrane. **A:** Cryo tomograms of U2OS cells reveal bell-shaped assemblies at the inner membrane. **1:** Central tomographic slice of a tomogram showing multiple convex structures at the mitochondrial inner membrane (white arrowheads). **2:** Central tomographic slice at higher magnification. The convex structures feature a hollow core (side view). **3:** Central tomographic slice at higher magnification. The structure appears as a ring (top view). **B:** Bell-shaped structures are a frequent feature of mammalian mitochondria. **1:** Central slice of an axonal mitochondrion of a rat hippocampal neuron. **2:** Central slice of a mitochondrion of a COS-7 cell. **C:** Quantification of convex complexes upon down regulation of PHB1 and PHB2. Boxes represent median, lower and upper quartiles. Whiskers represent SD. ***p < 0.001. **D:** Rendering of the prohibitin subtomogram average (U2OS cells, 2.5 Å pixel size) results in a round bell-shaped assembly. **1:** Side view. **2:** Top view indicating 11 units composing the bell-shaped prohibitin complex. **3:** Clipping of the side view average. The grey dashed lines indicate the location of the mitochondrial inner membrane (MIM). Scale bars: 200 nm (A1, B), 25 nm (A2, 3).

**Fig. 3.:**
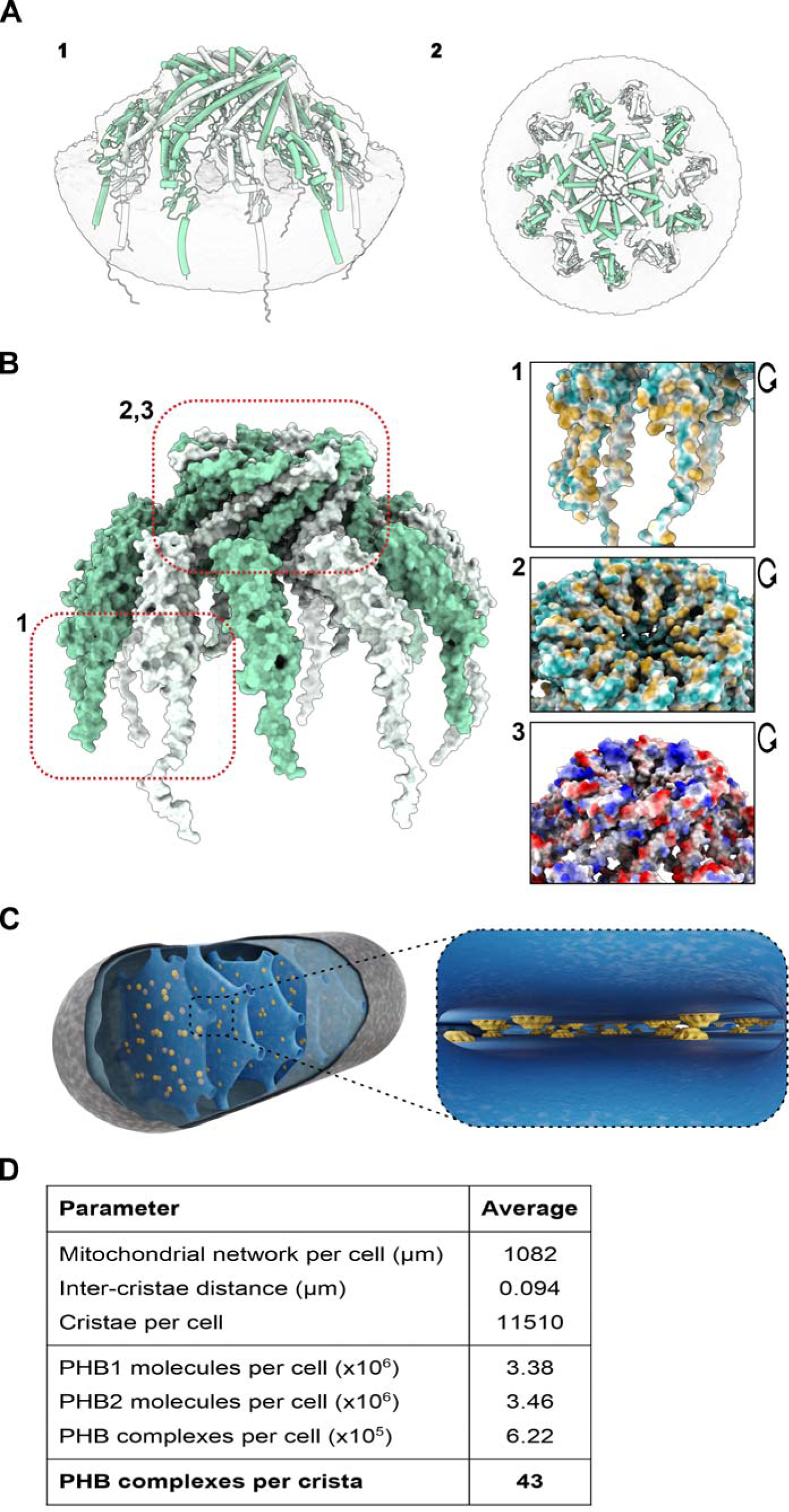
Molecular modelling of the human prohibitin complex. **A:** Cartoon representation. Predicted structures of PHB1 (green) and PHB2 (white) molecules were positioned into the final cryo-EM map (light grey, transparent). **1:** Side view. **2:** Top view. **B:** Surface representation (left). Interfaces of neighbouring prohibitin molecules (right). **1, 2:** Hydrophobicity surface. Orange colors represent hydrophobic side residues. Hydrophobic transmembrane domains anchor prohibitins to the membrane (1). The top of the prohibitin complex is stabilized by the hydrophobic C-termini (2). **3:** Electrostatic potential surface. Blue, red: Negatively and positively charges side chains, respectively. **C:** Cartoon representing the abundance and distribution of prohibitin complexes in a mitochondrion. Right: View into the crista lumen. **D:** Mitochondrial key numbers and prohibitin abundance.

To investigate whether endogenously expressed tagged PHB1 or PHB2 interacted with untagged prohibitins, co-immunoprecipitation experiments using nanobodies against DK were performed on mitochondria purified from knock-in cells (**Fig. S5**). When using PHB1-DK and PHB2-DK as bait proteins in co-immunoprecipitations, untagged PHB1 as well as PHB2 were pulled down. This indicates that the tagged prohibitins formed complexes with endogenous untagged PHB1 and PHB2.

To determine the distribution of PHB1-DK and PHB2-DK within the mitochondrial inner membrane, we performed quantitative immunogold electron microscopy (EM) on both knock-in cells lines^10, 11^. A highly specific antibody against DK was employed to label the target proteins. The localization of each gold particle associated with the mitochondria was assigned either to the inner boundary membrane (IBM) or the crista membrane (CM). For PHB1-DK and PHB2-DK we found 75.4% and 84.5% of the gold particles at the crista membranes, respectively (**Fig. 1C, D**). Despite considering that the ratio of IBM to CM was 0.64 to 1 in the U2OS cells, the concentration of the prohibitions is about three to five times higher in the CM than in the IBM (**Fig. S6A**). This finding was somewhat surprising, as in budding yeast the prohibitins are almost evenly distributed between crista membrane and inner boundary membrane^12^. To verify the predominant crista membrane localization of the prohibitins in U2OS cells, we performed immunogold EM using antibodies directed against PHB1 and PHB2. We found 75.4% and 81.4% of the gold particles to be localized in the CM, confirming the predominant localization of the prohibitins in the CM.

Crista resident proteins such as the respiratory chain complexes have been shown to exhibit a low mobility within the mitochondrial network^13–15^. To test if also PHB1 and PHB2 exhibit low mobility, fluorescence recovery after photobleaching (FRAP) experiments were performed, using the genome-edited cell lines expressing either PHB1-DK or PHB2-DK (**Fig. 1E, F**). We found recovery of both proteins to be very slow (in the minute range), with immobile fractions of 85% to 90%. In comparison, matrix-targeted DK exhibited sub-second recovery rates with an immobile fraction of < 10%. These findings are fully in line with the majority of the prohibitins being localized in the crista membranes.

Next, we aimed at determining the average number of PHB1 and PHB2 molecules per cell. To this end, recombinant His-tagged PHB1 and PHB2 proteins were purified from *Escherichia coli* and used as a reference in quantitative Western blotting (**Fig. S6B**). It was determined that on average each U2OS cell contains approximately 3.38 ± 0.23 10^6^ molecules of PHB1 and 3.46 ± 0.15 10^6^ molecules of PHB2. These numbers are in good agreement with previous mass spectrometry studies using various human cell lines that found between 3.1 and 9.8 10^6^ PHB1/PHB2 molecules per cell^16^. In budding yeast, prohibitins have been postulated to form ring-like complexes out of 12-20 PHB1/PHB2 dimers^3, 4^. For human cells, no direct evidence for prohibitin rings is available. Still, if we were to assume that human cells also contain rings of 16 prohibitin dimers a single U2OS cell would contain approximately ∼2.14 10^5^ prohibitin rings. As 75-84% of all PHB1/PHB2 are localized to the crista membranes, it could be estimated that on average about 1.7 10^5^ prohibitin rings formed by 16 dimers would be located on the crista membranes.

To estimate how many prohibitin rings might be located on a single crista, we next determined the average number of cristae in U2OS cells. These cells contain mostly stacked lamellar cristae. To determine their average number, the average total length of the mitochondrial network and the average number of cristae per micrometer mitochondrial tubule was determined using light and electron microscopy, respectively (**Fig. S6C, D**). The total mitochondrial network length of U2OS cells was 1082 ± 360 μm and the stacked crista membranes had a distance of 74 ± 15 nm. As groups of stacked lamellar cristae are separated by membrane voids^17^, which cover 21 ± 8% of the mitochondrial tubules, the corrected average distance between two crista membranes was 94 ± 19 nm (**Fig. S 6E, F**). Based on these measurements, an average of 11,510 cristae is estimated per U2OS cell.

Based on these data and the assumption that U2OS cells exhibit prohibitin rings consisting of 16 dimers, it can be calculated that each lamellar crista is expected to carry on average ∼15 prohibitin rings, so that each crista side would exhibit seven or eight prohibitin rings. Assuming a ring diameter of 20 nm, these rings would cover approximately 0.7% to 1.4% of the available crista surface^4^.

We reasoned that if in U2OS cells indeed about 1% of the cristae surface is covered by a sizeable ring-like structure, these structures should be visible in cryo-electron tomograms and potentially amenable to subtomogram averaging.

Consequently, we aimed to identify about 20 nm sized ring-shaped structures in the crista membranes of human cells. To this end, vitrified human U2OS cells were thinned into ∼150 nm thick lamellae using cryo-focused ion beam milling and then imaged with cryo-electron tomography (cryo-ET)^18^.

Within the mitochondrial inner membrane, we observed numerous convex structures. The majority of these prominent structures were localized in the crista membranes but were also localized in the inner boundary membrane (**Fig. 2A1**). In the top view, these structures appeared as a ring with a diameter of about 20 nm, whereas the side view revealed a convex shape with a height of about 9 nm, with the body of the structure pointing towards the inter membrane space, i.e. the crista lumen or the space between inner boundary membrane and outer membrane (**Fig. 2A2, 3, Fig. S7A**). We conclude that the size, the localization, as well as the inner membrane orientation of these structures fitted to the proposed elusive prohibitin structure.

As the prohibitins are well conserved, we suspected that similar structures should also exist in mitochondria of other cell types. Hence, we used cryo-ET to study tomograms of mitochondria of cultured rat hippocampal neurons and of COS-7 cells. These features were also present in both cell types, indicating a common structural element within the inner membranes of mammalian mitochondria (**Fig. 2B1, 2**).

To investigate whether the identified structures are indeed prohibitin complexes, we determined the abundance of these features in tomograms of wild-type cells, and cells downregulated for PHB1 or PHB2. siRNA mediated knockdown of either PHB1 or PHB2 led to the absence of both prohibitins (**Fig. S7B**, **C**)^19, 20^. First, we scrutinized the tomograms of PHB1 and PHB2 knock down (KD) cells for the convex structures and normalized their number to the mitochondrial volume. We found a strong reduction in the number of structures in both PHB1KD (246 ± 123 particles/µm^3^) and PHB2KD (199 ± 92 particles/µm^3^) cells compared to the abundance in wild type cells (686 ± 328 particles/µm^3^, **Fig. 2C**, **Fig. S7D**). Second, we investigated tomograms of U2OS PHB2-DK cells. On the convex structures in mitochondria of these cells, we frequently found additional densities close to the top that indicate the presence of DK molecules in these complexes (**Fig. S7E-G**). In comparison, we were unable to detect similar features in mitochondria of U2OS wt cells. We thus conclude that the convex structures are prohibitin complexes.

For detailed structural insights into prohibitin complexes, we employed subtomogram averaging to obtain an average 3D map from subvolumes containing these structures^21^. To this end, we manually picked 817 particles and aligned them along the z-axis followed by subsequent particle pose optimization in Dynamo (**Fig. S8A**)^22^. An alignment of two independent half maps resulted in a resolution of 28.1 Å for the initial average (0.143 criterion, 3.97 Å pixel size, **Fig. S8B**). This cryo-EM map revealed a bell-shaped structure that resembles a ring in the top view with a large empty cavity in the center. We proceeded to extract the CTF-corrected subtomograms in Warp for subsequent automated 3D refinement in Relion4.0 with no initial symmetry implied^23, 24^. Initially, we aligned all particles at 5 Å pixel size. The resulting cryo EM map revealed 11 densities in the top view that could represent subunits of the prohibitin complex (**Fig. S8C**).

Based on this refined cryo-EM map, we proceeded with automated 3D refinement of the dataset with C11 symmetry implied. The resulting map showed a sharper representation of the lipid bilayer and provided a clearer view on the protein bell. A Fourier shell correlation revealed a resolution of 18.3 Å for the cryo-EM map (0.143 criterion, pixel size 5 Å, **Fig. S8D, E**).

Finally, we re-extracted the subtomograms at 2.5 Å pixel size for automated 3D refinement with implied C11 symmetry, resulting in a final cryo-EM map of the prohibitin assembly at 16.3 Å resolution (FSC, 0.143 criterion, **Fig. 2D, Fig. S8F-H**). Subsequent 3D classification in Relion did not lead to further separation of the particles into distinct structural classes.

The final cryo-EM map revealed a circular bell-shaped structure that is embedded into the lipid bilayer. We measured a diameter of 190.2 Å directly above the lipid bilayer. The whole bell had a height of 84.0 Å measured from the lipid bilayer to the top of the particle and a narrowed top with a diameter of 89.0 Å. The particles were embedded into either the IBM or the CM. The opposing membrane thus was the mitochondrial outer membrane or the opposite CM of the crista. Notably, the map revealed a large empty cavity on top of the bilayer enclosed by the bell. Additionally, upon manual inspection of individual tomograms, no evidence was found to suggest occasional occupancy of this gap. This suggests that the gap is not occupied by large proteins by default.

Next, we aimed at modelling the molecular architecture of the prohibitin complex based on the final cryo-EM map. As no experimentally obtained structural data of prohibitins are available, we relied on AlphaFold2 to predict the structures of full length PHB1 and PHB2 (**Fig. S9A1, 2**). We placed these predicted structures manually into the experimentally obtained cryo-EM map.

Numerous possible arrangements were tested. We found that each of the 11 densities in the cryo-EM map fit one molecule of either PHB1 or PHB2 respectively. Accordingly, the entire prohibitin complex would consist of 11 molecules in total.

Ultimately, because biochemical evidence suggests alternating PHB1 and PHB2 molecules, successive PHB1 and PHB2 molecules were placed with the N-terminal transmembrane domains located in the lipid bilayer^3^. Thereby, the coiled-coil domains were oriented in perpendicular to the lipid bilayer. The final molecular model comprised 11 monomeric PHB molecules forming a bell-shaped assembly fitting into the cryo-EM map (**Fig. 3A1, 2**). In this model, the N-terminal domains of both, PHB1 and PHB2 are anchored in the lipid bilayer (**Fig. 3B1**). This positioning of the coiled coil domains seems to support only a weak interaction of individual prohibitin molecules with neighboring prohibitin molecules. The C-termini interact with each other and thereby stabilize the center of the bell-top. The PHB domains that form the top of the bell structure are additionally stabilized by alternating negative (PHB1:232-252, net. negative −3) and positive (PHB2: 244-264, net. positive +5) charged amino acid sidechains (**Fig. 3B2, 3**, **Fig. S9B1, 2**). Indeed, previous biochemical data showed that the strongest interaction of individual protein-subunits is between helical PHB domains of adjacent prohibitins^3, 25^.

To test the plausibility of the final model, we employed molecular dynamics (MD) simulations of the entire predicted complex in GROMACS (**Fig S9C, D**)^26^. The coiled-coil domains showed fluctuations relative to the position of the globular domains. These fluctuations included small tilts of the coiled-coil domain to the sides and are more prominent to the N-terminal part of the coiled-coil domain, which is in line with weaker densities in the cryo-EM map at these positions (**Fig. S9E**). This suggests that the N-terminal part including the coiled coil domain is intrinsically dynamic, while the C-terminal part of the PHB bell is more stable. The molecular model remained stable over 100 ns of MD simulation without major alterations, suggesting that it represents a plausible model for the molecular structure of the prohibitin complex in human cells.

The overall bell-shaped structure of the prohibitin complex resembled the structure formed by the bacterial prohibitin homologues HflK/C^5, 6^. The HflK/C structure associates with three hexameric complexes of proteases. Intriguingly, our final cryo-EM map did not reveal any further associated proteins even though the prohibitin complex has been demonstrated to be involved in the regulation of mitochondrial AAA proteases (mAAA protease)^27–29^. While we cannot rule out an occasional association with the mAAA protease, this interaction might play a minor role in U2OS cells.

The prohibitin complex has a C11 symmetry formed of monomers, whereas the HflK/C complex exhibits a C12 symmetry of dimers. Consequently, the bell-like structure formed by 12 HflK/C dimers has a height of ≥ 13 nm and is thereby substantially larger that the prohibitin structure (∼9 nm). As the mitochondrial intermembrane space is distinctly narrow (∼10 nm) the rather spacious HflK/C complex would not fit, whereas the smaller prohibitin complex has a suitable size (**Fig. S9F**). With its matching dimension that almost spans the space between the two membranes, the prohibitin complex might even be involved in tethering the two opposing membranes.

As the overall bell-shaped structure formed by the prohibitins resembles the structure formed by the HflK/C complex, this arrangement seems to be a core structural principle underlying the function of SPFH proteins^5, 6, 30^. Quantification of the number of cristae and the number of prohibitin complexes suggests that on average about 43 complexes are localized in a single crista (**Fig. 3C, D**). The fact that the bell-shaped prohibitin complexes occur with such a high frequency might open new strategies to explore the therapeutic potential of interfering with prohibitin function.

## Supporting information

Supplement Lange et al

## Acknowledgements

We thank Tanja Koenen, Nicole Molitor, Christian Dienemann and Ulrich Steuerwald for excellent technical assistance and Hauke Hillen for discussions. This work was supported by the Deutsche Forschungsgemeinschaft (DFG, German Research Foundation) under Germany’s Excellence Strategy - EXC 2067/1- 390729940 and by the European Research Council (ERCAdG No. 835102) (both to SJ). It was funded by the DFG-funded TRR 274 (to SJ), SFB 1456 (to SJ), FOR 2848 (project Z01 to DR and SJ).

## Author contributions

FL, MR, JND and PI performed experimental work, analysed data and prepared figures. DR and DW carried out electron microscopy experiments at room temperature. SJ supervised the study. FL, MR and SJ wrote the manuscript with input from all authors.

## Materials and methods

### Plasmids

#### Overexpression plasmids

Cloning of constructs for fusion protein overexpression was carried out using the primers listed in **Table S1**. The Dreiklang DNA sequence was amplified via PCR from the plasmid pQE31-Dreiklang^9^. PHB1 and PHB2 coding sequences were amplified from plasmids provided by the Human ORFeome clone collection with the internal IDs 6030, 394 and 56919, respectively. The PCR products were purified and used for one-step isothermal assembly with EcoRV-digested pFLAG-CMV-5.1 (Sigma, St. Louis, MO, USA)^32^.

#### Nuclease plasmids

Design of gRNAs was done using the CRISPR Design Tool (http://crispr.mit.edu). Annealed oligonucleotide pairs (**Table S2**) were cloned into the bicistronic vector pX330 as described in detail previously^33,34^.

#### Donor plasmids

DNA sequences for the left and right homology arms were amplified from genomic DNA using the primer pairs listed in **Table S3**. The Dreiklang DNA sequence was PCR-amplified from the plasmid pQE31-Dreiklang^9^. The PCR products were purified and cloned into EcoRV-digested pUC57 (Fisher Scientific, Schwerte, Germany) using one-step isothermal assembly followed by site-directed mutagenesis to remove Cas9 recognition sites^32^.

### Cell culture

U2OS and COS-7 cells (American Type Culture Collection, Manassas, VA, USA) were cultured in Dulbecco’s modified Eagle’s medium (DMEM) (Invitrogen, Carlsbad, CA, USA) supplemented with 10% fetal bovine serum (PAA, Pasching, Austria), 100 units/mL penicillin, 100 μg/mL streptomycin (all Biochrom, Berlin, Germany), and 1 mM sodium pyruvate (Sigma, St. Louis, MO, USA) under constant conditions at 37 °C and 5% CO_2_. Plasmid transfections were done using FuGENE HD (Promega, Mannheim, Germany) according to the manufacturer’s instructions. Mitochondria of living cells were stained using 100 nM MitoTracker Deep Red FM (Fisher Scientific, Schwerte Germany) at 37 °C for 30 min.

#### Cell culture for cryo electron microscopy

Confluent cultures of either U2OS wt or Cos7 wt cells were first detached by trypsination and subsequently suspended in DMEM complete medium. Next, 3 ml of the cell suspension was placed in ibidi glass bottom dishes and 4 coated grids were added to on the bottom of the dish. Cells were allowed to adhere to the grid surface for at least 18 hours prior to plunge freezing.

#### Hippocampus isolation and dissociation

Rat hippocampi were obtained through resection from Wistar rats. Newborn rats (P0 or P1) were decapitated, and the brain was carefully extracted to preserve its integrity. The isolated brain was placed in a 3.5 cm culture dish filled with ice-cold HBSS without calcium and magnesium (Thermo Fisher Scientific). After separating the hemispheres and removing meninges, the hippocampus was isolated, placed in a 15 ml Falcon tube, and stored on ice.

For dissociation, the hippocampi were placed in 4.5 ml ice-cold HBSS without calcium and magnesium, and 0.5 ml of freshly thawed 2.5% trypsin (Thermo Fisher Scientific) was added. Dissociation occurred for 18 minutes in a water bath set to 37 °C with occasional gentle shaking. Digestion was halted by adding 10 ml pre-warmed DMEM supplemented with 5% FBS to each Falcon tube, followed by centrifugation at 100 g for 3 minutes. After discarding the supernatant, the tissues were washed twice with 10 ml pre-warmed HBSS without calcium and magnesium.

Subsequently, tissues were dissociated using a 5 ml plastic Pasteur pipette in approximately 5 ml Neurobasal plating medium. The cell suspension was filtered through a 40 µm cell strainer into a 50 ml Falcon tube. The final cell suspension was achieved by diluting the strained cell suspension with pre-warmed Neurobasal plating medium to reach a final cell count of 1 10^6^ cells/ml. Cells were then plated by adding 1 ml of the final cell suspension onto coated Quantifoil grids. One hour after plating, the supernatant was replaced with standard cultivation medium. Hippocampal neurons were cultivated until 18 days in vitro (DiV) before freezing.

### Generation of knock-in cells

U2OS or HeLa cells were co-transfected with the respective combination of nuclease and donor plasmids for targeting either PHB1 or PHB2. Seven to ten days after transfection, the cells were subjected to single cell sorting into 96-well plates using a FACSAria II (BD Biosciences, Heidelberg, Germany). About two weeks after sorting, the cells of each well were split and equally distributed into two wells of a 24-well plate with one of the two wells containing a glass cover slip for microscopic inspection. Cells that expressed the fluorescent fusion protein were identified using an epifluorescence microscopy (DM6000B, Leica Microsystems, Wetzlar, Germany) equipped with an oil immersion objective (1.4 NA, 100x, Planapo, Leica) and a BGR filter cube (excitation: 495/15, emission: 530/30). Successfully targeted clones were expanded and genotyped via PCR using the primers listed in **Table S4**.

For DNA sequencing, genomic DNA was isolated from selected clones, the respective on- or off-target site was PCR amplified (**Tables S4-S5**) followed by purification and ligation into a pCR™Blunt II-TOPO® vector using a Zero Blunt® TOPO® Kit (Thermo Fisher Scientific, Waltham, MA, USA) according to the manufacturer’s instructions. Plasmids containing an insert were identified via colony PCR and 15 to 20 plasmids were sequenced per locus.

### siPool mediated knock-downs of prohibitins

We transfected U2OS cells with 3 nM of PHB1 or PHB2 siPools (siTOOLs Biotech, Germany), respectively following the manufacturer instructions. The cells were first cultivated for 2 days in standard 6-well plates. After 2 days, the cells were detached and the cell suspension transferred to PLL/fibronectin coated SiO_2_ R1/4 grids and allowed to adhere for 24 hours prior to plunge freezing in liquid ethane at melting point. The duration of siPool knock-downs for both prohibitins (PHB1 & PHB2) was 3 days in total.

### Immunoblotting

Cells were grown to 80-85% confluence and detached from the growth surface and counted using a Scepter™ 2.0 Cell counter (EMD Millipore, Billerica, MA, USA). Cells were harvested by centrifugation followed by lysis of 10^6^ cells in 100 μl of radioimmunoprecipitation assay (RIPA) buffer supplemented with 1 mM EDTA, 1 mM PMSF, 10 U/ml universal nuclease (Thermo Fisher Scientific, Waltham, MA, USA) and 1x complete protease inhibitor cocktail (Roche, Basel, Switzerland). After addition of RIPA buffer, the cell suspension was placed on ice for 30 min with vortexing steps every 10 min. The suspension was centrifuged at 13 000 rpm at 4 °C for 30 min. The supernatant was collected, and the protein concentration measured using the Pierce BCA protein assay kit (Thermo Fisher Scientific, Waltham, MA, USA). Samples were diluted to 1.2 µg/µl with RIPA buffer and mixed with the respective amount of 6x Laemmli buffer (375 mM Tris pH 6.8, 12% SDS, 60% glycerol, 0.6 M DTT, 0.06% bromophenol blue) to a final concentration of 1 µg/µl. The suspension was boiled at 95 °C for 5 min, flash frozen in liquid nitrogen and stored at −20 °C for further use.

Extracts corresponding to a known number of cells as well as recombinant 6xHis-PHB1 or 6xHis-PHB2 (Abcam, Cambridge, UK) were separated on 4-15% Mini-Protean^®^ TGX™ Precast Gels (Bio-Rad, Munich, Germany) according to the manufacturer’s instructions. Separated proteins were transferred to a nitrocellulose membrane (GE Healthcare, Freiburg, Germany) in transfer buffer (25 mM Tris, 190 mM glycine, 20% methanol) at 4 °C for 16 h. The membrane was rinsed in Tris-buffered saline (TBS) with 0.1% Tween-20 (TBST) and incubated in 5% blocking buffer (5 g skim milk per 100 mL TBST) at room temperature for 30 min. Primary antibodies were diluted in blocking buffer and incubated with the membrane at room temperature for 1 h. The following primary antibodies were used: anti-PHB1 (EP2803Y, 1:2000, Abcam, Cambridge, UK), anti-PHB2 (EPR14523, 1:5000, Abcam, Cambridge, UK), anti-GFP (JL-8, 1:3000, Clontech, Saint-Germain-en-Laye, France) and anti-Actin (AC74, 1:3000, Sigma-Aldrich, St. Louis, MO, USA). After washing with TBST, the membranes were incubated at room temperature for 1 h with HRP-conjugated anti-rabbit or anti-mouse secondary antibodies (Dianova, Hamburg, Germany) diluted 1:5000 in blocking buffer. After washing with TBST, the membrane was incubated with Pierce ECL western blotting substrate (Thermo Fisher Scientific, Waltham, MA, USA) and exposed to a CCD camera. Membranes were stripped by incubation with Restore™ (Thermo Fisher Scientific, Waltham, MA, USA) at 37 °C for 30 min followed by the described protocol for reprobing with a different antibody.

### Isolation of mitochondria and immunoprecipitation

Mitochondria were isolated from wildtype, PHB1-DK or PHB2-DK U2OS cells grown to 80-85% confluence, detached from the growth surface and harvested by centrifugation. The cell pellet was resuspended in THE buffer (300 mM trehalose, 10 mM 4-(2-hydroxyethyl)-1-piperazineethanesulfonic acid–KOH, pH 7.7, 10 mM KCl, and 1 mM EDTA) supplemented with 0.1% (w/v) bovine serum albumin (BSA) and lysed using a Dounce homogenizer. Homogenized cells were subjected to centrifugation at 11000 x g at 4 °C for 10 min. The mitochondria-containing pellet was resuspended in THE buffer without BSA and the protein concentration measured using the Bradford Protein Assay (Bio-Rad, Munich, Germany). The protein concentration was adjusted to 10 µg/µl with THE buffer.

GFP-Trap-Agarose beads (Chromo-Tek, Munich, Germany) were equilibrated in ice-cold dilution buffer (20 mM Tris-HCl pH 7.5, 50 mM NaCl, 0.5 mM EDTA, 10% (v/v) Glycerol, 0.3% (w/v) Digitonin, 1 mM PMSF) and mixed with 800 µg of mitochondrial extract. After rotation at 4 °C for 1 h, beads were centrifuged and washed ten times using dilution buffer. The beads were resuspended in 2x Laemmli buffer (125 mM Tris pH 6.8, 4% (w/v) SDS, 20% (v/v) glycerol, 0.2 M DTT, 0.02% (w/v) bromophenol blue), boiled at 95 °C and centrifuged. The supernatant was used for subsequent SDS-PAGE and western blot analysis.

### Fluorescence microscopy

#### Sample preparation

Cells were cultured on glass cover slips until they reached a confluence of about 70-85% and fixed in 37 °C prewarmed 4% (w/v) formaldehyde in PBS at RT for 5 min. The cells were permeabilized using 0.5% (v/v) Triton-X-100 in PBS for 5 min followed by subsequent incubation in blocking buffer (5% (w/v) BSA in PBS containing 100 mM glycin) for 15 min. Primary antibodies were diluted in blocking buffer and coverslips were incubated with that solution at room temperature for 1 h. The following primary antibodies were used: rabbit anti-PHB1 (EP2803Y, 1:250, Abcam, Cambridge, UK), rabbit anti-PHB2 (EPR14523, 1:500, Abcam, Cambridge, UK); mouse anti-ESR1 (D12, 1:500, Santa Cruz Biotechnology), rabbit anti-GFP (ab290, 1:1000, Abcam, Cambridge, UK). After three washing steps in PBS, fluorophore-coupled secondary antibodies were diluted 1:1000 and added for incubation at room temperature for 1 h. The following secondary antibodies were used: sheep anti-mouse and goat anti-rabbit (all Dianova, Hamburg, Germany) coupled to KK114^35^ or Alexa 594 (Atto-Tec, Siegen, Germany). After three PBS washing steps, cells were embedded in Mowiol 4-88 mounting medium containing 1 µg/ml 4′,6-Diamidin-2-phenylindol (DAPI) and 2.5% (w/v) 1,4-diazabicyclo-[2,2,2]-octane (DABCO).

#### Confocal microscopy

Confocal imaging was done using the Leica TCS SP8 Confocal Microscope (Leica, Wetzlar, Germany). All recordings were performed using a pinhole diameter of one Airy unit (1.22λ/NA), a scan speed of 400 Hz and a 63x oil immersion objective (HCX PL APO CS 63x/1.40-0.60 oil). The following laser lines were used for fluorescence excitation: a 405 Diode (405 nm), an argon laser (458 nm/ 476 nm/ 488 nm/ 496 nm/ 514 nm) and a helium-neon (HeNe) laser (633 nm). Fluorescence detection was done using photomultipliers (PMTs) and a Hybrid Detector (HyD) operated within the dynamic range. Separation of excitation and emission light was accomplished using an AOTF (acousto-optic tunable filter). Multicolor imaging was done using sequential acquisition between frames. For image digitization a sampling rate according to the Nyquist criterion was chosen. Each image was recorded at least twice for averaging.

#### FRAP analysis

FRAP measurements were done using a Leica TCS SP5 Confocal Microscope and the FRAP Wizard application (Leica, Wetzlar, Germany). All recordings were performed using an open pinhole and a 63x oil immersion objective (HCX PL APO CS 63x/1.40-0.60 oil). Living U2OS cells were mounted in a custom-built live-cell chamber and maintained at 37 °C in CO_2_-independent Leibovitz L-15 medium (Thermo Fisher Scientific, Waltham, MA, USA) for imaging.

Mitochondria of U2OS cells expressing mitochondrial matrix targeted Dreiklang (mito-DK) were photobleached in a circular region of interest (ROI) with a diameter of about 0.5 µm using an argon laser at 20% laser power. Imaging after bleaching was performed at a time interval of 19 msec for about 2 sec with the 514-nm line of the argon laser. Mitochondria of U2OS knock-in cells expressing PHB1-DK or PHB2-DK were bleached in a ROI with a diameter of about 2.5 µm using an argon laser at 20% laser power. Imaging after bleaching was performed at a time interval of 8 sec for about 1 min with the 514-nm line of the argon laser.

### Electron microscopy

#### Plastic embedding

U2OS cells were grown on Aclar^®^ polymer cover slips until 80-85% confluence. Cells were prefixed in 2.5% (w/v) glutaraldehyde in 0.1 M sodium cacodylate (pH 7.4) at RT for 15 min postfixed in the same buffer at 4 °C for 15 h. Cells were washed three times in 0.1 M sodium cacodylate (pH 7.4) and incubated in 1% (w/v) OsO_4_ in 0.1 M sodium cacodylate (pH 7.4) for 3 h. Cells were washed once in 0.1 M sodium cacodylate (pH 7.4) and then twice in water. The cells were place in 0.1% (w/v) uranyl acetate (in H_2_O) for 30 min. Uranyl acetate was washed out by subjecting the cells to 30% ethanol three times for 5 min followed by dehydration through a 50%, 70% and 100% ethanol series. Afterwards the cells were placed in 100% propylene oxide for 5 min and then transferred to 50%/50% propylene oxide/Epon for 1h followed by placement to 100% Epon overnight. Samples were sectioned to 50 nm thickness with a Leica EM UC6 ultramicrotome (Leica EM UC6, Leica Microsystems, Wetzlar, Germany). Each section was transferred to 0.7% (w/v) Pioloform coated 200 mesh carbon grids. Samples were subjected to post contrasting using 1% (w/v) uranyl- and lead acetate. Electron microscopic recordings were acquired using a Philips CM 120 transmission electron microscope equipped with a TVIPS 2k x 2k slow-scan CCD camera (Philips, Amsterdam, Netherlands).

#### Immunogold labelling

U2OS cells were grown to 80-85% confluence and fixed in 37 °C prewarmed 4% (w/v) PFA (paraformaldehyde) in PBS at RT for 30 min. Further sample processing was done according to previous work^11^. Samples were sectioned into 80 nm thin slices and incubated with diluted primary antibodies for 30 min. The following antibodies were used: anti-GFP (JL-8; 1:20, Clontech, Saint-Germain-en-Laye, France), anti-PHB1 (EP2803Y, 1:20, Abcam), anti-PHB2 (EPR14523, 1:40, Abcam).

Subsequently each sample was incubated with protein A coupled to 10 nm gold particles for 20 min followed by multiple washing steps and additional contrasting using uranyl acetate/methylcellulose on ice for 10 min. Electron microscopic recordings were done using a Philips CM 120 transmission electron microscope equipped with a TVIPS 2k x 2k slow-scan CCD camera (Philips, Amsterdam, Netherlands).

### Determination of crista area occupied by PHB1/2 complexes

First, we determined that the inner diameter of an average mitochondrial tubule in human U2OS cells is 564 nm (**Fig. S6 G**). Thus, the area of a circular crista membrane (A = π r²) is around 249,832 nm^2^. Each lamellar crista consists of two opposing membranes hence the total membrane area per crista is around 499,664 nm^2^. Second, we assumed a ring-like PHB complex with a diameter of about 20 nm which corresponds to an area of about 314 nm^2^ per PHB ring^2, 4^. Thus, a single PHB ring occupies 0.063% of the total available crista area while 11 to 22 PHB rings per crista occupy between 0.7% and 1.4% of the available crista surface, respectively.

### Cryo Electron Microscopy

#### Grid preparation for cell culture

Quantifoil R2/1 or SiO_2_ R1/4 gold grids (both Quantifoil Mirco Tools GmbH, Germany) were briefly washed in chloroform and placed on drops of poly-D-lysine (100 µg/ml). After incubation for 1 hr at room temperature the grids were washed three times on drops of HBSS for 10 min each. Subsequently, the grids were placed on drops of fibronectin (20 µg/ml) and incubated for 1 hr at room temperature.

Finally, the grids were washed three times on drops of HBSS for 10 minutes each and directly used for cell plating.

#### Preparation of frozen-hydrated cells

Grids containing cells were picked up with a pair of tweezers and any residual medium was removed by manual blotting at the tweezer tips. The sample was then rapidly loaded into a Vitrobot (ThermoFisher, Germany) set at 37°C and 95% humidity. We set a wait time of 120 seconds and applied 3 µl of a solution containing 10% glycerol in HBSS (ThermoFisher, Germany). The grids were then backside blotted for 30 seconds with blot force 20 and subsequently plunged into liquid ethane at melting point. The grids were kept at liquid nitrogen temperatures until further processing.

#### Automated preparation of cryo-lamellae

Cryo-lamellae preparation was automated using an Aquilos 2 cryo FIB-SEM (ThermoFisher, Germany). Grids were initially coated with GIS platinum for 20 seconds. Lamella positions were identified around the center marker of the grids after automated tile acquisition with Maps (200x magnification, 5 kV, 13 pA). These positions were then transferred to Cryo AutoTEM with a target lamella size of 14 µm length and approximately 170 nm thickness, with a 4 nm offset after final polishing, resulting in lamellae of approximately 180 nm thickness. Following the automated procedure, the lamellae were additionally sputter-coated with metallic platinum (30 mA, 1 kV, for 1 to 2 seconds). The final lamellae were stored at liquid nitrogen temperatures until further imaging.

#### Cryo electron tomography and tomogram reconstruction

Cryo electron tomograms were recorded on a Titan Krios G2 microscope (ThermoFisher, Germany) operated at 300 kV, equipped with a FEG, BioQuantum imaging filter (Gatan, Germany) with a 20LeV energy width, and a K3 direct electron detector (Gatan, Germany). For automated acquisition of the tomograms we used SerialEM^36^. Recorded frames were motion corrected using Warp and dose filtered prior to tomogram reconstruction. Aligned frames from Warp were utilized for patch tracking or fiducial-based alignment of small platinum particles in IMOD for tomogram reconstruction. CTF correction was performed through phase flipping in IMOD. Tomograms for visualization and IsoNet training were reconstructed with binning 6 (11.90 Å pixel size) and an additional SIRT-like filter in

IMOD, set to 50 iterations. A comprehensive overview of all acquired tomograms used in this study is provided in **Table S6**.

#### Missing wedge correction in IsoNet

For missing wedge correction in IsoNet, we employed the tool at binning 6 (11.90 Å pixel size) for visualization and PHB quantifications^37^. IsoNet was trained separately using 4 tomograms from U2OS wt cells, 4 from Co7 wt cells, and 4 from rat hippocampal neurons. CTF deconvolution was not applied before masking. A mask was generated with density and std percentages set at 60, and a z-crop of 0.05. Subtomograms (120 per input tomogram) were extracted, and the network underwent training for 35 iterations with 15 epochs each. Noise was introduced at iterations 16, 21, 26, and 31 with levels of 0.05, 0.1, 0.15, and 0.2, respectively. The resulting model was used for predicting tomograms depicted in this study.

#### Subtomogram averaging in Dynamo

For subtomogram averaging, 37 tomograms from U2OS wt cells were utilized, and 817 particles were manually picked in Dynamo at a pixel size of 3.87 Å^38^. An initial average was created by manually aligning 50 particles in dgallery. Subsequently, particles were globally aligned to the initial average using Dynamo’s “global search” pre-set. Randomization of particle orientations was performed using the “dynamo_table_randomize_azimuth” function, and the resulting table was used for particle averaging. Pose-optimized particles were extracted in Warp at a 5 Å pixel size and aligned in Relion4.0 through 3D auto-refinement, revealing an 11-fold symmetry of the PHB complex. All particles were then aligned to a C11-fold symmetrized template. The resulting star file from auto-refinement was used to re-extract particles in Warp at a 2.5 Å pixel size, followed by another round of 3D auto-refinement in Relion. Resolution estimation for all alignments was performed using the EMDB FSC server with two independent halfmaps generated from the Relion4.0 3D auto-refinement (https://www.ebi.ac.uk/emdb/validation/fsc/results/). Visualization of 3D models of the cryo-EM maps was conducted using ChimeraX^39^.

### Modelling

For the modelling of the PHB complex, we used the Alphafold predicted structures of PHB1 (Uniprot P35232) and PHB2 (Uniprot P50093) respectively. As starting point, the structure of Prohibitin 1 was choosen^40^. After placement of the globular domain in the density map, backbone rotations between residues 174-178, 229-233 were made, analogous to the bacterial homologous complex, while a C11 symmetry was maintained using PyMOL^6, 41^. Afterwards, the resulted model was fitted into the cryo-EM derived density map using the molecular dynamics method from Flex-EM, with a cap-shift of 0.5 Å, 20 runs and 100 iterations per run^42^. Additionally, all secondary structure elements from the Alphafold model with high confidence (plDDT>70), were treated as rigid bodies. This provided the C-terminal structured part of the interacting helices (residues 176-253), while a new placement of more globular and transmembrane part (residues 1-174) was made, since due to their membrane interaction, a fitting in a density map including membranes is not anticipated. Using this arrangement, a homology model of an alternating complex of 5 PHB1 and 6 PHB2 was built using MODELLER 10.4^43^. Stereochemical restraints were added as well as helical restrains on the transmembrane domain for PHB2 (residues 19-39). To avoid interference of the longer disordered C- and N-terminal part of PHB2, these areas were redirected during modelling by setting a Z-coordinate upper limit for residues 1-19 of 107 Å with a standard deviation of 0.3 Å and a lower limit for residues 286-291 of 158 Å with the same standard deviation. The coordinates frame was identical as the cryo-EM derived map. From 300 models, after ranked by DOPE score, two were selected and visual inspected^44^. All-atom systems including solvent and a lipid membrane were built using CHARMM-GUI as described previously^45, 46^. The membrane consisted of POPC lipids in a hexagonal setup. After equilibration, 100 ns of a molecular dynamics simulation were computed for each setup using GROMACS 2021.3 and the CHARMM36m force filed as described before^45, 47, 48^. Density maps were computed using GROmaps using a pixel size of 2.5 Å^49^.

### Blender modelling of an average mitochondrion

The mitochondrion was drawn to scale in Blender 4.0.2 using the following data: length, 1.5 µm; outer membrane diameter, 565 nm; inner membrane diameter, 540 nm; distance outer and inner membrane, 25 nm; crista diameter (without junctions), 480 nm; crista thickness, 25 nm (for clarity reasons); inter-cristae distance, 175 nm (for clarity reasons); thickness of all membranes, 5 nm.

### Quantification and statistical analysis

For the data shown in Fig. 3C we used the Jarque-Bera test to assess the normal distribution of the data followed by the two-tailed unpaired t-test (T) to determine *p*-values. These results are summarized in **Table S7**.

