## Supplement Lange et al for "In-situ architecture of the human prohibitin complex"

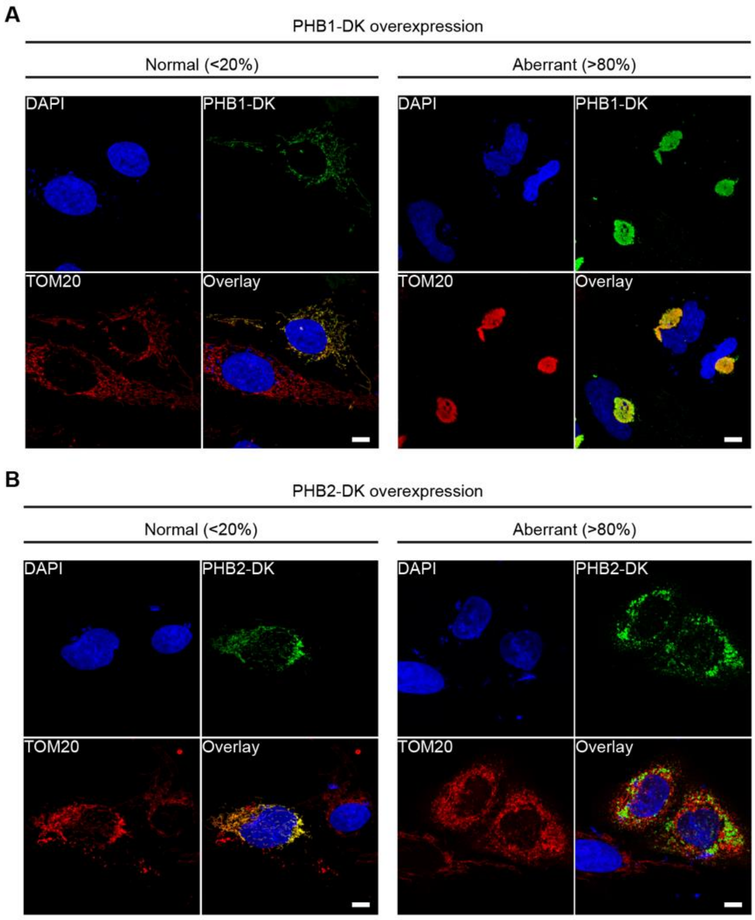

**Fig. S1: Overexpression of PHB1-DK and PHB2-DK in human cells**. U2OS cells were transfected with a plasmid encoding PHB1-DK (**A**) or PHB2-DK (**B**). For each construct, three separate transfection experiments were performed; each time more than 100 fluorescent cells were analysed, and mitochondrial morphology was classified as “normal” or “aberrant”. Scale bars, 10 µm

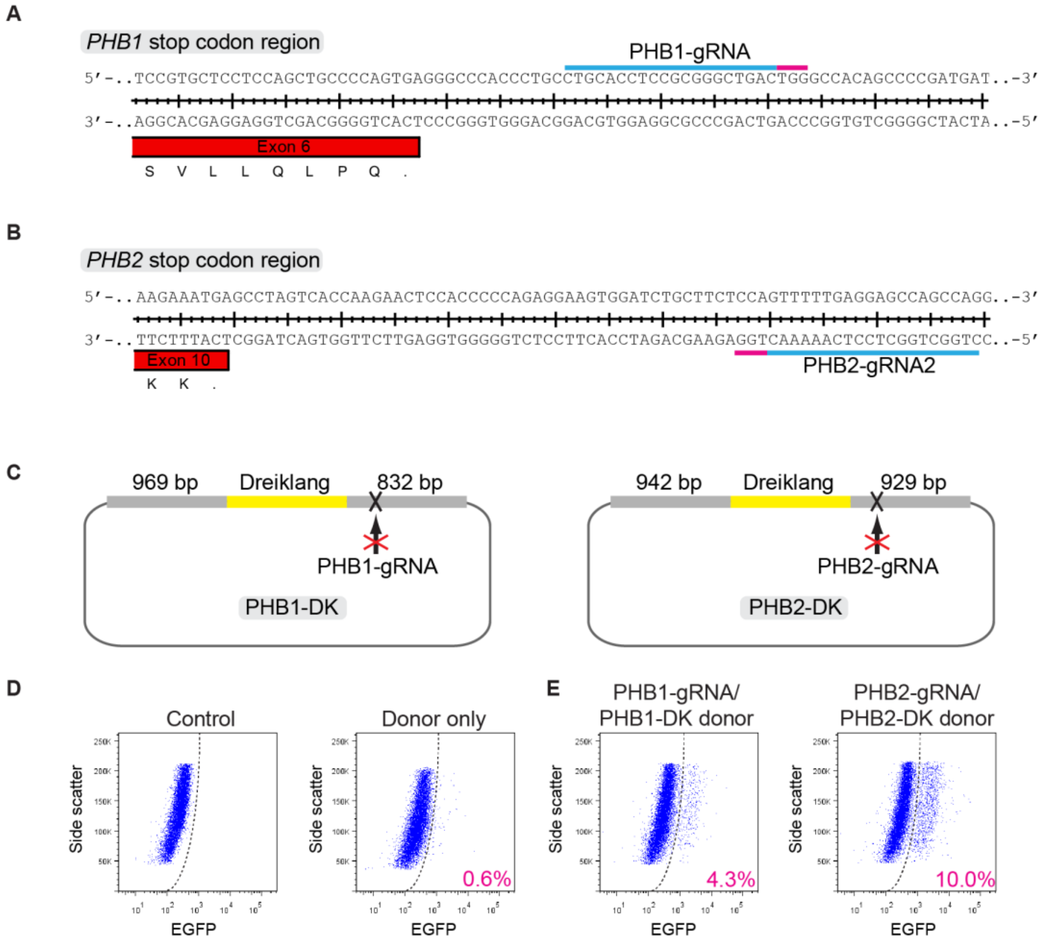

**Fig. S2: Guide RNAs and donor plasmids for C-terminal tagging of human prohibitins**. (**A, B**) Two gRNAs were designed for targeting the stop codon region of human PHB1 (**A**) or PHB2 (**B**), respectively. Light blue: gRNA binding site; Magenta, protospacer adjacent motif (PAM site. (**C**) Donor plasmids encoding the fluorescent protein Dreiklang (DK) flanked by gene-specific homology arms containing point mutations in the PAM sites rendering the respective construct resistant to Cas9-mediated degradation. (**D, E**) FACS analysis of U2OS cells. Untransfected (control) and donor plasmid only (donor only) transfected U2OS cells were used as a negative control to set FACS gates (**D**). Co-transfection of PHB1-gRNA1 or PHB2-gRNA2 with Cas9-resistant PHB1-DK or PHB2-DK donor plasmids (**E**). The mean fraction of DK+ cells of three independent experiments is shown in magenta.

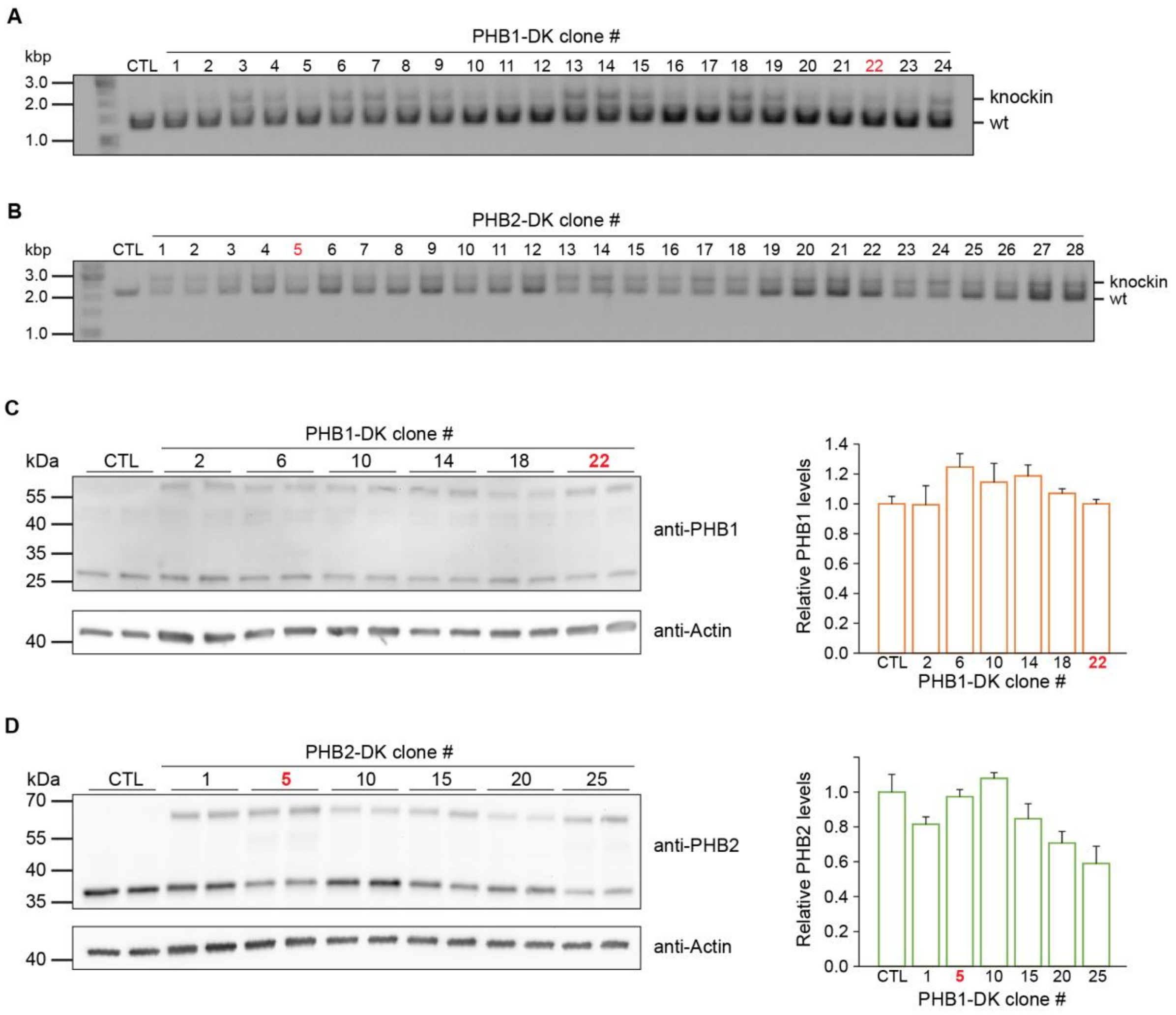

**Fig. S3: Genotyping and expression levels in PHB1-DK and PHB2-DK knock-in clones**. (**A, B**) PCR analysis of 24 PHB1-DK clones (**A**) and 28 PHB2-DK clones (**B**) obtained after single cell selection using FACS and clonal expansion. CTL, parental, non-edited control cell line. (**C, D**) Protein expression level analysis in five PHB1-DK (**C**) and five PHB2-DK clones (**D**). Whole cell extracts of clones were analyzed via immunoblotting using antibodies against PHB1 (**C**) or PHB2 (**D**), respectively. Wildtype extract (CTL) was loaded as a reference and actin was detected as internal loading control. Band intensities were quantified, PHB1 (**C**) or PHB2 (**D**) expression levels corrected for variations in loaded amounts and normalized to the PHB1 (**C**) or PHB2 (**D**) expression level in wildtype (CTL) cells. Clone numbers highlighted in red indicate the respective clone chosen for subsequent analysis.

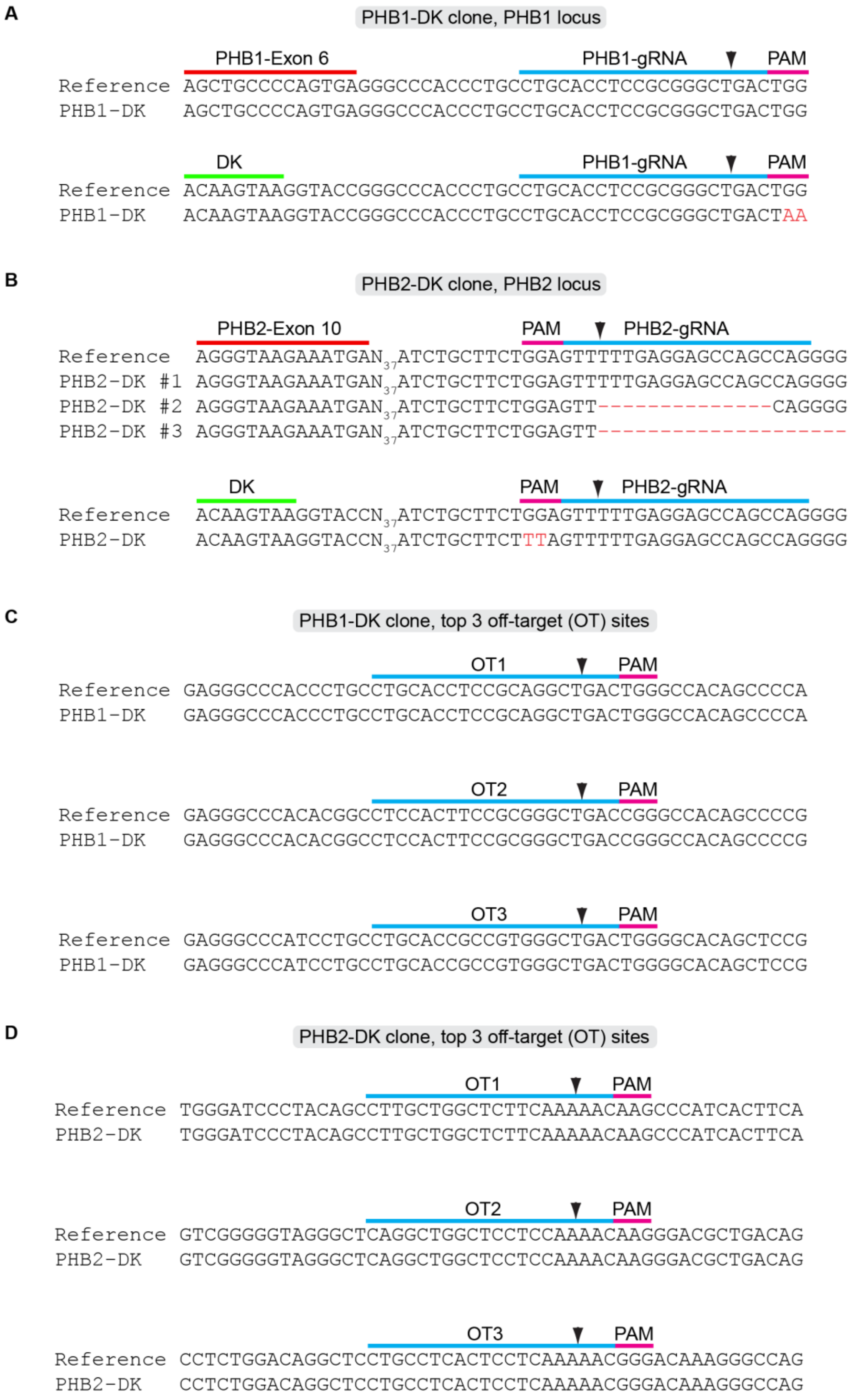

**Fig. S4: Sanger sequencing of on-target and predicted possible off-target sites in PHB1-DK and PHB2-DK knock-in clones**. (**A**) Sanger sequencing of the selected heterozygous PHB1-DK clone revealed that the untagged PHB1 allele is unmodified (top) and that the tagged

PHB1-DK allele contains the expected mutations introduced into the donor plasmid (bottom). (**B**) Sequencing of the selected heterozygous PHB2-DK clone revealed that the untagged PHB2 alleles contain three DNA sequence variants (top): unmodified sequence (#1), a 14 bp deletion (#2) or a 20 bp deletion (#3). The tagged PHB2-DK allele contains the expected mutations introduced into the donor plasmid (bottom). (**C, D**) Absence of mutations at the predicted top three possible off-target sites for the respective gRNA used to generate PHB1-DK (**C**) and PHB2-DK (**D**) suggests precise gene editing. For all experiments, the respective DNA regions were amplified from genomic DNA, subcloned and sequenced using Sanger sequencing at a depth of at least 3x.

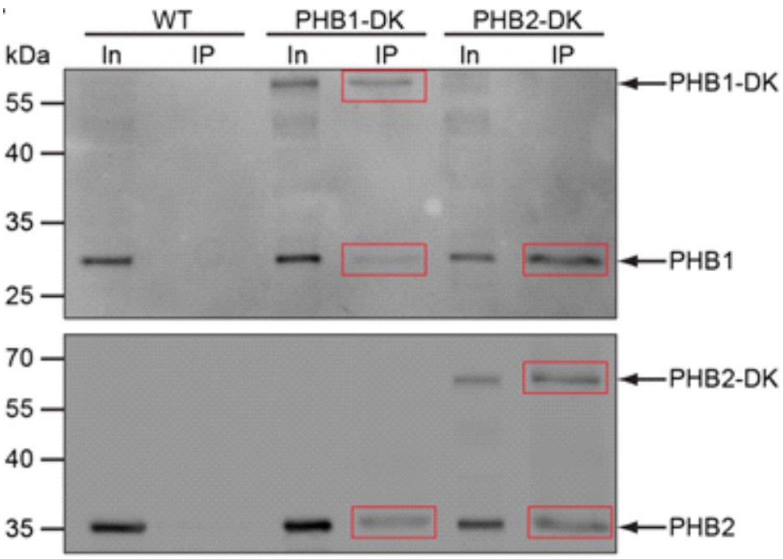

**Fig. S5: Co-immunoprecipitation of prohibitins.** Co-immunoprecipitation of prohibitins from isolated mitochondria. Mitochondrial lysates from PHB-DK1 or PHB-DK2 knock-in cells were immunoprecipitated using Dreiklang as the bait protein. Antibody detection; upper blot: PHB1, lower blot: PHB2.

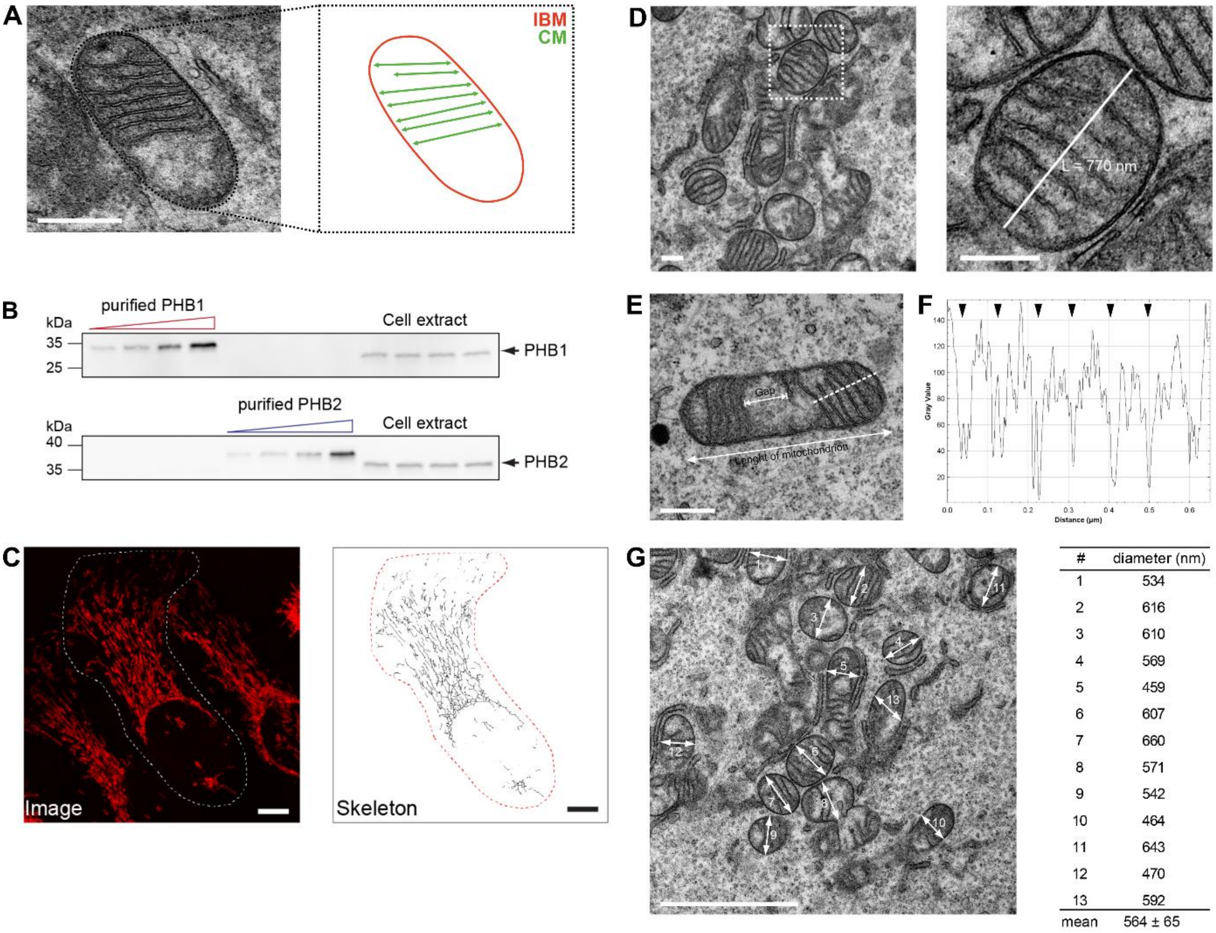

| # | diameter (nm) |
|---|---|
| 1 | 534 |
| 2 | 616 |
| 3 | 610 |
| 4 | 569 |
| 5 | 459 |
| 6 | 607 |
| 7 | 660 |
| 8 | 571 |
| 9 | 542 |
| 10 | 464 |
| 11 | 643 |
| 12 | 470 |
| 13 | 592 |
| mean | 564 ± 65 |

**Fig. S6: Quantitative analysis of PHB complexes.** (**A**) Representative TEM micrograph of a U2OS mitochondrion. The IBM to CM ratio was determined by measuring the total IBM length (red) and summing the lengths of all cristae, multiplied by 2 due to the dual-membrane composition of a crista. The U2OS wt cells exhibited an IBM to CM ratio of 0.64 to 1 (n = 32 mitochondria). (**B**) Quantitative analysis of prohibitin abundance in cells. Increasing amounts of purified PHB1 and PHB2 were used as a reference to estimate the total number of PHB molecules iin U2OS cells. Purified prohibitins and U2OS cell extracts were loaded on the same gel and blotted for antibody-based detection of PHB1 (top) or PHB2 (bottom). Note that purified PHB1 and PHB2 run at a higher molecular weight than endogenous PHBs due to the His-tag and linker sequence. (**C**) Determination of total mitochondrial network length by image analysis. U2OS cells stained with Mitotracker Deep Red FM revealed a mitochondria network length per cell of 1082 ± 360 µm (mean ± SD, n = 34 cells). A representative image is shown with a network length of 1029 µm. (**D**) Representative TEM micrograph of a U2OS mitochondrion. Average distance between crista membranes determined from individual mitochondria was determined as 74.1 ± 15.1 nm, SD, n = 36 mitochondria. (**E**) Representative TEM micrograph of a U2OS mitochondrion used to determine the total mitochondrial length and size of voids between groups of crista. Measurements revealed an average gap size of 21 ± 8% of the mitochondrial tubules (n = 98 mitochondria). (**F**) Line profile along the white dashed line in (E). This mitochondrion shows an average inter-crista distance of approximately 79 nm. Cristae are indicated with black arrowheads. (**G**) Representative EM micrograph of U2OS cells. Data were used to measure the diameter of mitochondria. Scale bar in A, 500 nm. Scale bar in C, 10 µm. Scale bar in D, E 300 nm. Scale bar in G, 2 µm.

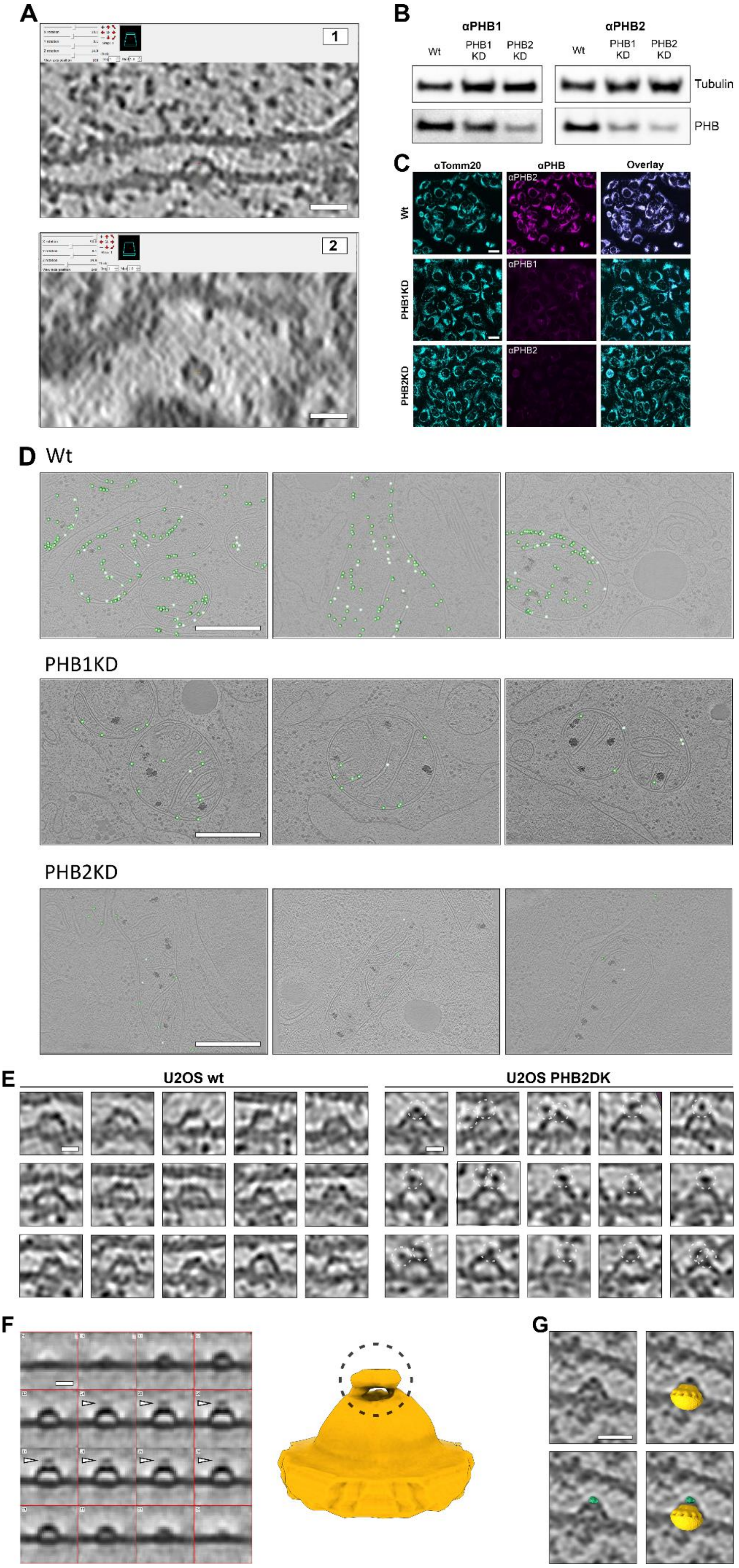
A
1
2
B
αPHB1
αPHB2
Wt
PHB1 KD
PHB2 KD
Tubulin
PHB
C
αTomm20
αPHB
Overlay
Wt
PHB1KD
PHB2KD
αPHB2
αPHB1
D
Wt
PHB1KD
PHB2KD
E
U2OS wt
U2OS PHB2DK
F
G

**Fig. S7: Prohibitin structures**. (**A**) IMOD slicer view of a representative particle recorded on mitochondria of wildtype U2OS cells. **1:** Side view shows a convex profile of the particle, tilted to 10° (x-rotation). **2:** Top view exhibits a ring-like appearance of the same particle, tilted to 90° (x-rotation) with respect to (1). (**B**) Western blot analysis. RNAi-mediated knockdown of PHB1 and PHB2 showed that depletion of one prohibitin leads also to the absence of the other (see also Fig. S10). (**C**) Immunofluorescence imaging of PHB1KD and PHB2KD cells using an antibody against PHB1 and PHB2, respectively. Mitochondrial network labelled with Tomm20 antibodies (cyan). (**D**) Representative central tomographic slices of U2OS cells. Prohibitin structures were marked with green spheres. PHB1KD and PHB2KD cells exhibit reduced prohibitin abundances compared to wt cells. (**E**) Central slices of tomograms of individual prohibitin complexes. PHB2-DK cells show additional densities at the top of the structures, indicating PHB2 molecules fused to Dreiklang. (**F**) Left: Subtomogram average of PHB-DK particles (U2OS PHB2-DK cells). STA confirms the presence of an additional density at the bell top, indicating the presence of DK (white arrowheads). Right: Isosurface rendering in ChimeraX highlights the additional density. (**G**) Central tomographic slice of a prohibitin assembly. Image overlay with a volume representation of the prohibitin cryo-EM map (orange) and a Dreiklang barrel (green). The additional density at the bell top matches the size of the Dreiklang barrel. Scale bars in A, 20 nm. Scale bars in C, 50 µm. Scale bars in D, 500 nm. Scale bars in E, F, 10 nm. Scale bar in G, 20 nm.

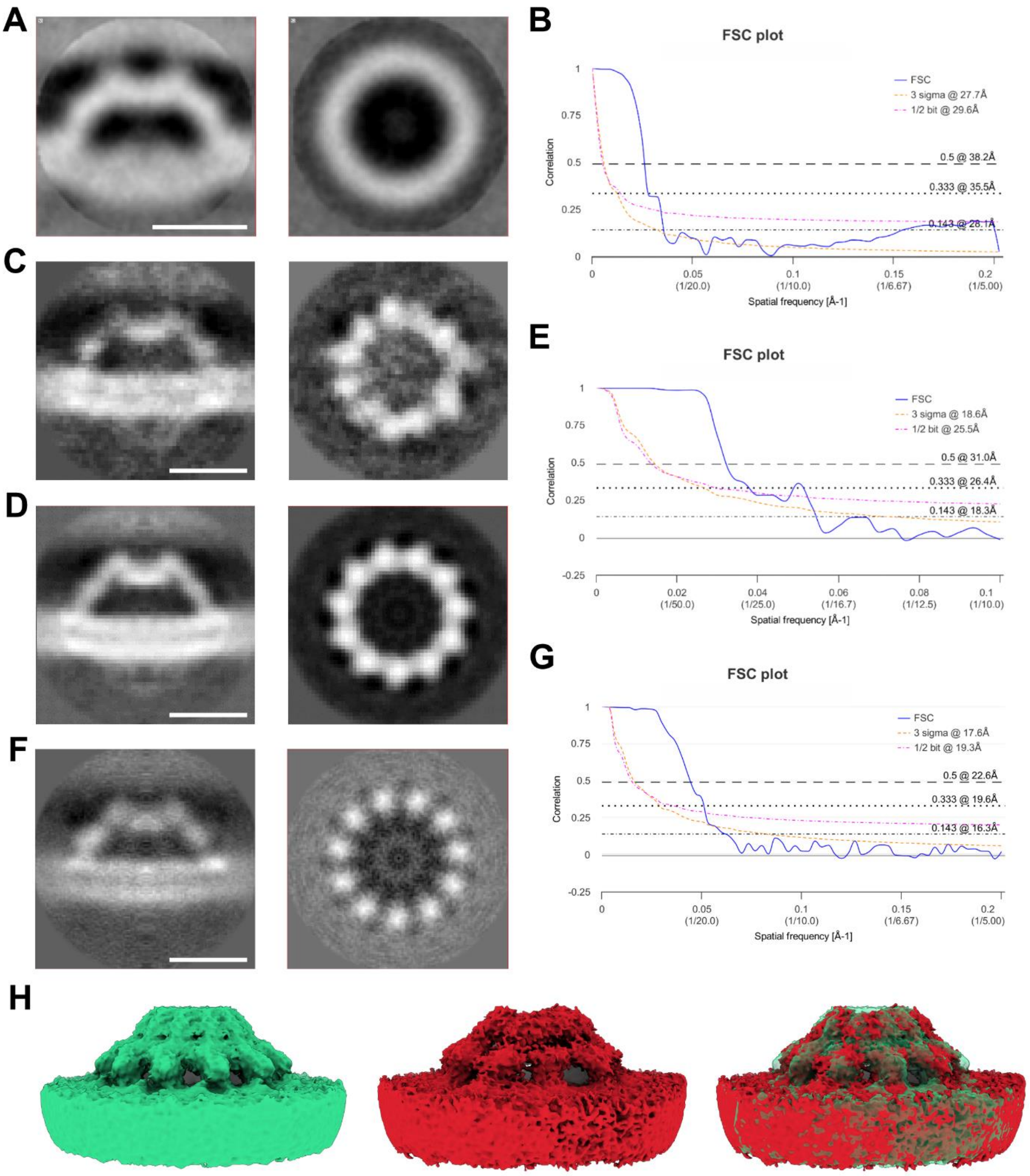

**Fig. S8: Subtomogram averaging of prohibitin complexes**. (**A**) Initial average of the PHB complex obtained through subtomogram averaging in Dynamo (3.87 Å pixel size). Left: X-view of the Cryo-EM map displaying a convex structure with a hollow core. Right: Z-view showing a ring-like structure. (**B**) Fourier shell correlation. A resolution of 28.1 Å (0.143 criterion) was obtained for the average presented in (A). (**C**) Average after initial alignment of subvolumes with 5 Å pixel size. The cryo-EM map suggests an underlying C11 symmetry of the PHB complex. (**D**) Subtomogram average of volumes with 5 Å pixel size aligned with C11 symmetry. Left: X-view of the average. Right: Top-view of the average. (**E**) Fourier shell correlation. A resolution of 18.3 Å (0.143 criterion) was obtained for the average presented in (D). (**F**)

Subtomogram average of volumes with 2.5 Å pixel size aligned with C11 symmetry. Left: X-view of the average. Right: Top-view of the average. (**G**) Fourier shell correlation. A resolution of 16.3 Å (0.143 criterion) was obtained for the average presented in (F). (**H**) Isosurface representations. The final cryo-EM map with C11 symmetry (green), alignment without symmetry (red), and an overlay of both maps. Scale bars, 10 nm.

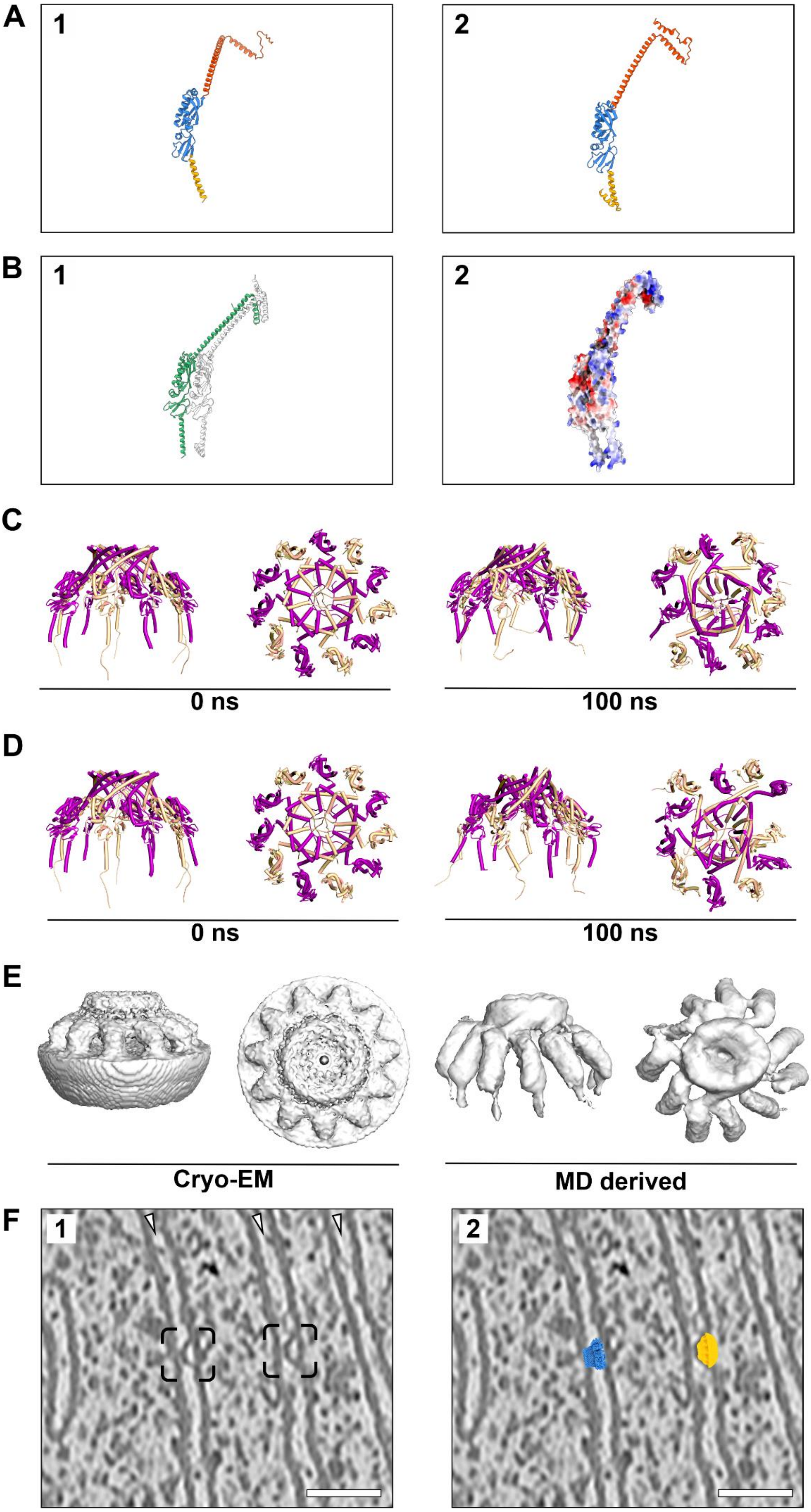

A
1
2
B
1
2
C
0 ns
100 ns
D
0 ns
100 ns
E
Cryo-EM
MD derived
F
1
2

**Fig. S9: Molecular diffusion of the PHB complex model**. (**A**) **1-2:** AlphaFold-predicted structures of the human prohibitins. Prohibitin 1 (PHB1, Uniprot P35232) and prohibitin 2 (PHB2, Uniprot Q99623) showing key domains – transmembrane (yellow), coiled-coil (blue), and PHB (orange). (**B**) **1-2:** AlphaFold-predicted model of the PHB1 (green) and PHB2 (white) dimer. Structural model (1). Electrostatic potential map (2). (**C**, **D**) Two models for the prohibitin complex used in molecular dynamics (MD) simulations (PHB1 in magenta, PHB2 in green). Preservation of the complex after 100 ns simulations is observed, with increased stability at individual PHB domains and flexibility at N-terminus and coiled-coil domains. (**E**) Isosurface representations. Comparison of the cryo-EM-derived map and an MD-derived map after 100 ns, excluding lipids for a protein-only comparison. The MD model maintains structural similarities with the initial model, indicating overall preservation during MD. (**F**) Size comparison of the bacterial HflK/C structure (blue) and the prohibitin complex (yellow) at mitochondrial membranes. **1:** Cryo electron micrograph of a U2OS mitochondrion. White arrowheads indicate crista lumen, and black boxes show the location of two prohibitin complexes. **2:** Structures fitted into the indicated locations, emphasizing that the bacterial HflK/C complex with 24 molecules (12 dimers) is too large for the crista lumen. Scale bar in E, 50 nm.

**Table S1: Primers used for Gibson assembly of overexpression plasmid**. All constructs are based on a pFLAG-CMV-5.1 backbone. DK, Dreiklang.

| Plasmid name | Primer | Sequence (5' to 3') |
|---|---|---|
| **PHB1-DK** | MR1143_PHB1_F | TTCATCGATAGATCTGATGCCACCATGGCTGCCAAAGTGTTTGAGTCC |
| | MR1144_PHB1_R | CCACTACCCTGGGGCAGCTGGAGGAGCAC |
| | MR1145_DK_F | TGCCCCAGGGTAGTGGTTCAGGGGTGAGCAAGGGCGAGGAGCTG |
| | MR1146_DK_R | GTCGACTGGTACCGATTTACTTGTACAGCTCGTCCATG |
| **PHB2-DK** | MR1147_PHB2_F | TTCATCGATAGATCTGATATGGCCCAGAACTTGAAGGACTTG |
| | MR1148_PHB2_R | CCACTACCTTTCTTACCCTTGATGAGGCTGTC |
| | MR1145_DK_F | TGCCCCAGGGTAGTGGTTCAGGGGTGAGCAAGGGCGAGGAGCTG |
| | MR1149_DK_R | GGTAAGAAAGGTAGTGGTTCAGGGGTGAGCAAGGGCGAGGAGCTG |
| **ESR1** | MR1216_ESR1_F | TTCATCGATAGATCTGATGCCACCATGACCATGACCCTCCACACC |
| | MR1217_ESR1_R | GTCGACTGGTACCGATTTAGACCGTGGCAGGGAAAC |

**Table S2: Primers used for gRNA cloning into pX330.**

| Gene | Primer | Sequence (5’ to 3’) |
|---|---|---|
| **PHB1** | MR716_F | CACCGCTGCACCTCCGCGGGCTGAC |
| | MR717_R | AAACGTCAGCCCGCGGAGGTGCAGC |
| **PHB2** | MR720_F | CACCGCTGGCTGGCTCCTCAAAAAC |
| | MR721_R | AAACGTTTTTGAGGAGCCAGCCAGC |

**Table S3: Primers used for Gibson assembly of donor plasmids**. All donor plasmids are based on a pUC57 backbone. LHA, left homology arm; RHA, right homology arm; DK, Dreiklang; SDM, site-directed mutagenesis.

| Donor plasmid | Primer name | Sequence (5’ to 3’) |
|---|---|---|
| **PHB1-DK** | GA265_LHA_F | TCTCGCGAATGCATCTAGATAGGTCACACGTTGCAGAGAGCTGTCTTCC |
| | GA266_LHA_R | ATCCGCTGCCCTGGGGCAGCTGGAGGAGCACGGAC |
| | GA267_DK_F | GCTGCCCCAGGGCAGCGGATCCGGCGTGAGCAAGGGCGAGGAGC |
| | GA268_DK_R | GGGTGGGCCCGGTACCTTACTTGTACAGCTCGTCCATG |
| | GA269_RHA_F | GTAAGGTACCGGGCCCACCCTGCCTGCACCTCCGC |
| | GA270_RHA_R | AGGCCTCTGCAGTCGACGATCCAGGAACGTAGGTCGGACACGTCTTTGGC |
| **PHB2-DK** | GA271_LHA_F | TCTCGCGAATGCATCTAGATATGTGTTACTCATTGCTGCACCCCT |
| | GA272_LHA_R | ATCCGCTGCCTTTCTTACCCTTGATGAGGCTGTCAC |
| | GA273_DK_F | GGGTAAGAAAGGCAGCGGATCCGGCGTGAGCAAGGGCGAGGAGC |
| | GA274_DK_R | GTGACTATGCGGTACCTTACTTGTACAGCTCGTCCATG |
| | GA275_RHA_F | GTAAGGTACCGCATAGTCACCAAGGACTCCACCCC |
| | GA276_RHA_R | AGGCCTCTGCAGTCGACGATGGTGCCTGAGATTCGGGAAGGCCTG |
| **PHB1-DK-PAM-mutagenesis** | MR1038_SDM | GCACCTCCGCGGGCTGACTAAACCACAGCCCC |
| | MR1039_SDM | GGGGCTGTGGTTTAGTCAGCCCGCGGAGGTGC |
| **PHB2-DK-PAM-mutagenesis** | MR1040_SDM | CAGAGGAAGTGGATCTGCTTCTTTAGTTTTTGAGGAGCCA |
| | MR1041_SDM | TGGCTCCTCAAAAACTAAAGAAGCAGATCCACTTCCTCTG |

**Table S4: Primers used for on-target site analysis via PCR and DNA sequencing.**

| Gene | Primer | Sequence (5’ to 3’) |
|---|---|---|
| **PHB1** | MR1269_in_F | CCTGCAGCCAACAGAGATGTATCCCTCC |
| | MR1051_out_R | GCAGTTTATACACATTTGTTTCCTTCCCAG |
| **PHB2** | MR1052_out_F | TTCCCTGACCTTTTGTTCTAATCATAGCTG |
| | MR1053_out_R | TCTAGTTGTTTATCTCATCTTAGCCTCCCA |

**Table S5: Primers used for off-target site analysis via DNA sequencing.**

| Gene | Primer | Sequence (5' to 3') |
|---|---|---|
| **PHB1** | MR1364_OT1_F | GAGATGGGGTCTCATCATGTTGCTCAGAC |
| | MR1365_OT1_R | CACAGAAGCAGTGGAAGCCAAACAGGTG |
| | MR1366_OT2_F | CACTTCGAACTGATGAGTCCTCAAACTGTAACC |
| | MR1367_OT2_R | GAAGGAGTTCACAGAAGCGATGGAAGCC |
| | MR1368_OT3_F | CGGGAAGGAGTTCACAGAAGCGGTGG |
| | MR1369_OT3_R | GCATGTGTCCTCATCCGCAAATCTTCCATC |
| **PHB2** | MR1370_OT1_F | CACAACTTGGTTTAGAATAAAGCCTCCACAGAC |
| | MR1371_OT1_R | CTTTTGACTTTTTAGGTTGACTTCCAGCCATG |
| | MR1372_OT2_F | CCGCATACATATTACCCACAATTCCCTTTCC |
| | MR1373_OT2_R | CATTCTGCCGTTTGTTACTTACCAATGCC |
| | MR1374_OT3_F | GAGACAGAGTCTTACTTTGTCACCCAGGCTG |
| | MR1375_OT3_R | GAATGAAGAAGGAAAAGTCTCTCATTCATTGCATC |

**Table S6: Recorded tomograms used in this study.** For each experiment, the sum of all recorded tomograms is listed. Each grid for each experiment is an independent sample preparation (grid preparation, cell batch, day of cell seeding and sample freezing) and thus, represents a biological replicate.

| Experiment | ∑ of tomograms | No. of grids | No. of biological replicates | pixel size (Å) | original magnification |
|---|---|---|---|---|---|
| Rat hippocampal neurons | 54 | 2 | 2 | 2.75 | 33000x |
| Cos7 wt cells | 27 | 2 | 2 | 2.75 | 33000x |
| U2OS wt cells | 92 | 4 | 4 | 1.98 | 42000x |
| U2OS PHB2DK | 65 | 3 | 3 | 1.98 | 42000x |
| U2OS wt cells | 73 | 4 | 4 | 2.75 | 33000x |
| U2OS PHB1 knock down | 41 | 3 | 3 | 2.75 | 33000x |
| U2OS PHB2 knock down | 41 | 3 | 3 | 2.75 | 33000x |

| Tomography collection and processing data | |
|---|---|
| Microscope | Titan Krios G2 |
| Voltage (kV) | 300 |
| Detector | Gatan K3 |
| Energy filter | yes (20 eV) |
| Total electron exposure per tilt series ($e^-/Å^2$) | approx. 120 |
| Defocus range | -2.5 to -4 |
| Tilt range (min, max, increment) | -60°, +60°, 3 |
| Tilt scheme | Dose-symmetric |
| Frame number | 3 |
| Tomograms used for STA (no.) | 37 |
| Initial subtomograms (no.) | 817 |
| Final subtomograms (no.) | 817 |
| Map resolution (Å at 0.143 FSC threshold) | 16.3 Å |
| Map resolution range | - |

**Table S7: Statistical tests for the abundance determination of prohibitin complexes in PHB1KD and PHB2KD cells.** We first used the Jarque-Bera test to assess the normal distribution of the data sets. The results indicated a normal distribution for all three samples and thus, we used a two-tailed t-Test to determine the p-values for all samples.

| **Jarque-Bera Tests** | | | | | | | | | | | |
|---|---|---|---|---|---|---|---|---|---|---|---|
| **Wt** | | | | **PHB1KD** | | | | **PHB2KD** | | | |
| observations | | n | 20 | observations | | n | 20 | observations | | n | 20 |
| sample skewness | | s | 0.820801 | sample skewness | | s | 1.057652 | sample skewness | | s | 0.538822 |
| sample kurtosis | | c | -0.50837 | sample kurtosis | | c | 0.691109 | sample kurtosis | | c | 1.338402 |
| JB test statistic | | JB | 2.46108 | JB test statistic | | JB | 4.126788 | JB test statistic | | JB | 2.460529 |
| **p-value** | | p | **0.292135** | **p-value** | | p | **0.127022** | **p-value** | | p | **0.292215** |

| t-Test: Two-Sample Assuming Equal Variances | | |
|---|---|---|
| | *Wt* | *PHB1KD* |
| Mean | 742.59 | 257.2864 |
| Variance | 113177.37 | 15886.57 |
| Observations | 20 | 20 |
| Pooled Variance | 64531.97 | |
| Hypothesized Mean Difference | 0.05 | |
| df | 38 | |
| t Stat | 6.04065606 | |
| P(T<=t) one-tail | 2.50335E-07 | |
| t Critical one-tail | 1.68595446 | |
| **P(T<=t) two-tail** | **5.0067E-07** | |
| t Critical two-tail | 2.024394164 | |

| t-Test: Two-Sample Assuming Equal Variances | | |
|---|---|---|
| | *Wt* | *PHB2KD* |
| Mean | 742.59 | 201.8522 |
| Variance | 113177.37 | 8877.388 |
| Observations | 20 | 20 |
| Pooled Variance | 61027.37965 | |
| Hypothesized Mean Difference | 0.05 | |
| df | 38 | |
| t Stat | 6.921285185 | |
| P(T<=t) one-tail | 1.57165E-08 | |
| t Critical one-tail | 1.68595446 | |
| **P(T<=t) two-tail** | **3.1433E-08** | |
| t Critical two-tail | 2.024394164 | |
